# SIRT6 is a key regulator of mitochondrial function in the brain

**DOI:** 10.1101/2022.11.08.515572

**Authors:** Dmitrii Smirnov, Ekaterina Eremenko, Daniel Stein, Shai Kaluski, Weronika Jasinska, Claudia Consentino, Barbara Martinez-Pastor, Yariv Brotman, Raul Mostoslavsky, Ekaterina Khrameeva, Debra Toiber

## Abstract

SIRT6 is implicated in DNA repair, telomere maintenance, glucose and lipid metabolism and, importantly, it has critical roles in the brain ranging from its development to neurodegeneration. In this work, we combined transcriptomics and metabolomics approaches to characterize the functions of SIRT6 in mice brains. Our analysis revealed that SIRT6 is a critical regulator of mitochondrial activity in the brain. In its absence, there is a mitochondrial deficiency with a global downregulation of mitochondria-related genes and pronounced changes in metabolites content. We predict that SIRT6 can affect mitochondrial functions through its interaction with the transcription factor YY1 that, together, regulate mitochondrial gene expression. Moreover, SIRT6 target genes include SIRT3 and SIRT4, which are significantly downregulated in SIRT6-deficient brains. Our results demonstrate that the lack of SIRT6 leads to decreased mitochondrial gene expression and metabolomic changes of TCA cycle byproducts, including increased ROS production, reduced mitochondrial number, and impaired membrane potential that can be partially rescued by restoring SIRT3 and 4 levels. Importantly, the changes observed in SIRT6 deficient brains are observed in brains of aging people, but the overlapping is greater in patients with Alzheimer’s, Parkinson’s, Huntington’s, and Amyotrophic lateral sclerosis disease. Overall, our results suggest that reduced levels of SIRT6 in the aging brain and neurodegeneration could initiate mitochondrial dysfunction by altering gene expression, ROS production and mitochondrial decay.

## Introduction

Aging is a consequence of the dysregulation of various self-maintenance mechanisms of a living system. Aging at the cellular level is accompanied by genomic instability, telomere shortening, loss of proteostasis, and mitochondrial dysfunction, together with a decrease in the efficiency of the DNA repair mechanism^1–3^. Moreover, these factors are interconnected. For example, the shortening of telomeres can lead to mitochondrial dysfunction^4^. Aging involves significant changes in the brain structure and functional capabilities^2,5–7^. Cognitive decline occurs naturally during aging, but in some cases, it can become pathological, such as in neurodegenerative diseases. Importantly, about 95% of neurodegenerative cases are age-related with no known genetic mutation. Therefore, a better understanding of the aging process in disease development is needed.

Sirtuins are a family of proteins that have mono-ADP ribosyltransferase or deacetylase activity^8–10^. As one of the most notable proteins of this family, SIRT6 is implicated in genomic stability^11–14^, DNA repair^13,15^, telomere maintenance^16^, cellular metabolism^17^ and importantly, it has critical roles in the protection against aging-associated diseases^18–20^ SIRT6-deficient mice have a progeroid-like syndrome with low body weight and a very short lifespan (~four weeks) compared to normal mice^11^. SIRT6 is widely expressed in mammalian brain tissues, with the highest expression level in cortical and hippocampal regions^21^. SIRT6 plays a neuroprotective role, protecting against DNA damage accumulation and during ischemic brain injury^18,22^. The lack of SIRT6, specifically in the brain, results in learning and memory impairments, increased DNA damage, and the promotion of cortical apoptotic cells, partially through the hyperphosphorylation and hyperacetylation of Tau^18,23^. In addition, through the changes in gene expression in these brains, we identified signatures of pathological aging brains, particularly relevant for A.D. and P.D., that could be partially reversed by calorie restriction^24^. Importantly, SIRT6 levels are decreased in the aging brains^18^ and even more pronounced in Alzheimer’s patients^23^, suggesting its involvement in age-related neurodegeneration and making it a good model to find the molecular mechanism of pathological aging in the brain.

One of the hallmarks of aging is mitochondrial activity impairment. Mitochondrion are vital cell organelles with many functions, including adenosine triphosphate (ATP) synthesis, calcium homeostasis handling, and lipid metabolism. ATP production occurs on the inner mitochondrial membrane, which incorporates five specific protein complexes (complexes I-V), forming an electron transport chain. The mammalian mitochondrial protein synthesis system involves genes from both nuclear and mitochondrial genomes. While mtDNA encodes only a small fraction of mitochondrial genes compared to nuclear DNA (1%), they are all necessary for synthesizing the respiratory complex proteins. To generate energy, electrons are transported through complexes I-IV moving across an electrochemical gradient to the ultimate acceptor, oxygen. This process is called oxidative phosphorylation (OXPHOS). As part of ATP production, various metabolites are formed in the mitochondria, such as *Acetyl-CoA, Citric Acid, Oxoglutaric acid, Succinic Acid, Malate*, and *Fumarate*. These metabolites control mitochondrial bioenergetics, and their altered levels might result in the deregulation of several aging-related pathways (e.g., mTOR, AMPK), implicating mitochondrial bioenergetic defects in aging^25–27^. During oxidative phosphorylation, the mitochondria also generates reactive oxygen species (ROS) molecules as a byproduct of ATP synthesis^28^. These molecules induce damage, which accumulates throughout the organismal lifespan and becomes harmful at high concentrations, inducing oxidative stress, DNA damage, and lipid peroxidation^29^. Since mtDNA is located near the ROS production sites, it might be more sensitive to oxidative damage and prone to possible mutations. The brain is particularly vulnerable to age-related mitochondrial damage because of its high energy demand^30^. Age-related accumulation of mitochondrial abnormalities disrupts synaptic transmission and neuronal metabolism, leading to neurodegeneration^31,32^. However, despite the clear role of mitochondrial dysfunction as a key marker of aging and neurodegenerative diseases, the exact mechanisms initiating this dysfunction are still poorly understood.

In this study, we emphasize the importance of SIRT6 for the essential mitochondrial processes in the brain, including oxidative phosphorylation and aerobic respiration. By using transcriptomics and metabolomics profiles of control and brain-specific SIRT6-deficient mice, we demonstrate a reduction in the expression of OXPHOS-related genes, as well as the abundance of tricarboxylic acid cycle (TCA) metabolites. We also validated these findings by measuring the mitochondrial membrane potential and mitochondrial content changes. To establish the regulatory mechanisms by which SIRT6 affects mitochondria, we focus on YY1, SIRT3 and 4 proteins. YY1 transcription factor is implicated in the regulation of mitochondria-related genes in skeletal muscle^33^ and has shown to share many cellular functions with SIRT6, including those related to neurodegeneration^24^. In turn, SIRT3 is reported to be a key deacetylase in the mitochondria, targeting OXPHOS, TCA cycle, and mitochondrial dynamics^34^. Because of these abilities, SIRT3 can contribute to the protection against oxidative stress, preventing neuronal cell death^35^. Finally, we link transcriptional changes of mitochondria-related genes with normal and pathological brain aging.

## Results

### Lack of SIRT6 alters gene expression levels in mouse brain

Brains missing SIRT6 functionality might present changes at multiple levels of molecular organization, from gene expression to metabolism. To explore these changes, we performed transcriptome profiling (RNA-seq) in brains derived from Wild Type (W.T., 4 samples) and brain-specific SIRT6-knockout (brSIRT6-KO, 4 samples) mice (Supplementary Tables 1, 2). In addition, we applied GC-MS and LC-MS techniques to quantify the abundance of 34 metabolites in W.T. (4 samples) and SIRT6-KO (4 samples) replicates derived from the SH-SY5Y cell line and complemented them with mouse Embryonic Stem Cell (mESC, 3 samples per W.T. and SIRT6-KO groups) metabolomics data. Then, we conducted a multilayer bioinformatics analysis of W.T. and SIRT6-KO transcriptomic and metabolomic profiles (Fig. 1a).

**Figure 1:**
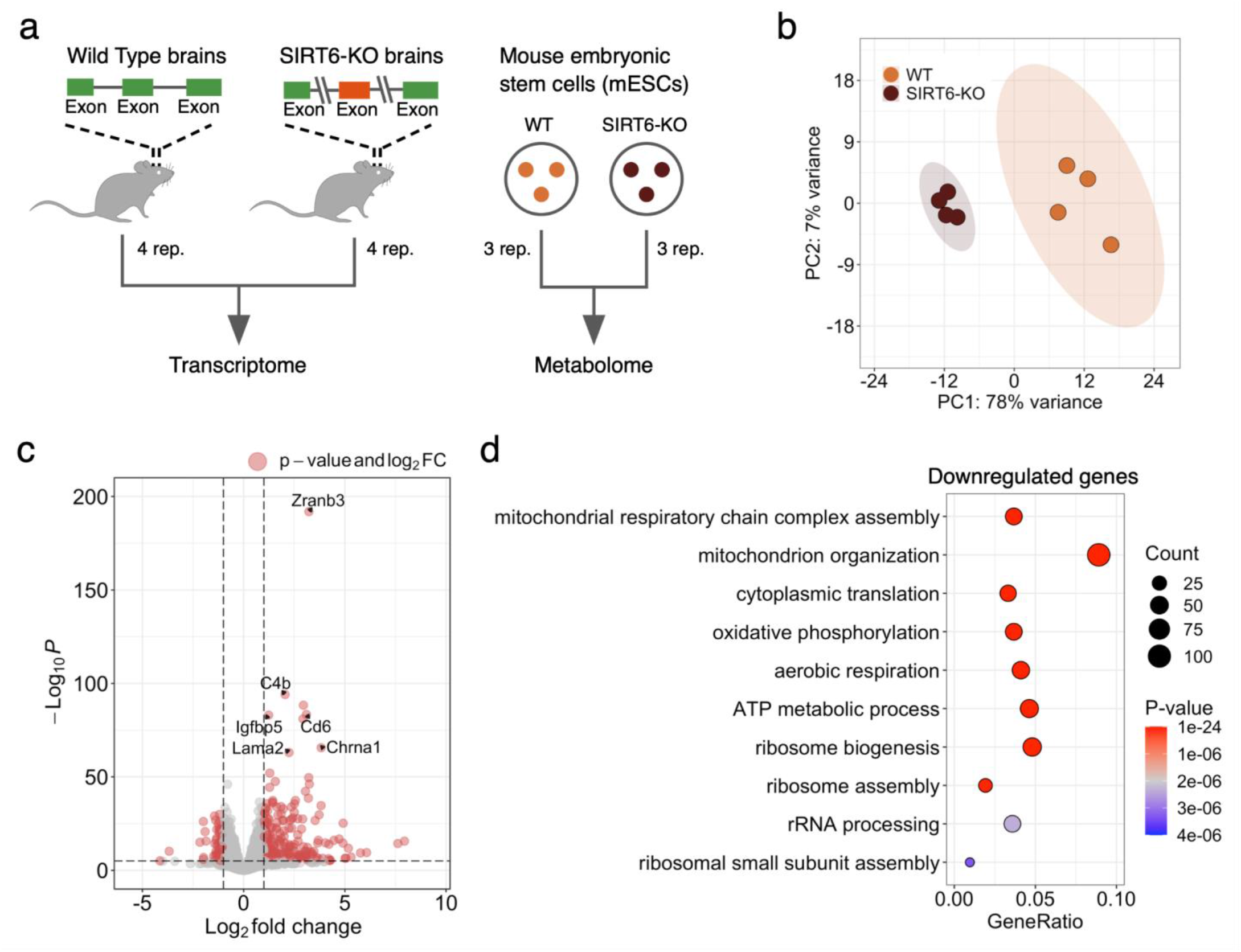
SIRT6 regulates gene expression levels. **(a)** Schematic illustration of the experimental design used in this study. Transcriptome profiles were collected from Wild Type (W.T.) and SIRT6 knockout (brSIRT6-KO) mouse brain samples. W.T. and brSIRT6-KO SH-SY5Y metabolomic profiles were complemented with metabolomics data on mouse embryonic stem cells. **(b)** Principal Component Analysis (PCA) plot showing separation between W.T. (orange) and brSIRT6-KO (brown) samples. Orange and brown ovals represent confidence ellipses of W.T. and brSIRT6-KO groups. **(c)** The volcano plot shows up and downregulated differentially expressed genes in brSIRT6-KO mice compared to W.T. mice. Red dots indicate significantly changed genes, and gray dots represent insignificant genes. **(d)** G.O. analysis showing the top 10 enriched biological processes for downregulated genes. Each circle corresponds to the enriched G.O. term and varies in size according to the number of significant genes belonging to this term. The gene ratio represents the number of D.E. genes belonging to the enrichment categories divided by the total number of genes per category.

Principal Component Analysis (PCA) of transcriptomic profiles revealed significant changes in gene expression levels between brSIRT6-KO and W.T. samples with a clear separation by the first principal component explaining 78% of the total variance (Fig. 1b). At the same time, transcriptomic profiles exhibited a high level of intra-group similarity, showing the Pearson’s R > 0.9986 for the W.T. group and Pearson’s R > 0.9992 for brSIRT6-KO replicates. In contrast, the inter-group Pearson’s R did not exceed 0.9978 (Supplementary Fig. S1a). Differential expression analysis between W.T. and brSIRT6-KO resulted in 2870 differentially expressed (D.E.) genes, ~85% of which were annotated as protein-coding sequences (Supplementary Table 3, Supplementary Fig. S1b). Consistent with the expected impaired deacetylase activity of SIRT6 upon knockout, 1481 DE genes exhibited elevated expression levels in brSIRT6-KO samples, while 1389 genes were downregulated (Fig. 1c). The list of top 10 significant features was represented exclusively by upregulated genes, including *Zranb3* (FDR p-value < 1.32×10^-192^), *C4b* (FDR p-value < 8.8×10^-95^), *Cd6* (FDR p-value < 4.18×10^-84^), as well as *Chrna1* (FDR p-value < 2.47×10^-66^) and *Lama2* (FDR p-value < 9.95×10^-64^) (Supplementary Fig. S1c), which were previously found among the most significant signatures of SIRT6 deficiency in the brains of the full-body K.O^24^. These results collectively indicate that SIRT6 deficiency has a major effect on transcriptional regulation in the mouse brain.

We further examined the functional roles of significant D.E. genes. G.O. enrichment analysis on upregulated genes revealed enriched terms associated particularly with *‘external encapsulating structure organization’* (FDR p-value < 3.7×10^-08^), *‘axon guidance*, and *‘neuron projection guidance’* (FDR p-value < 5.42×10^-08^ for both terms) (Supplementary Fig. 1d). Conversely, the list of downregulated features in W.T. compared to brSIRT6-KO was significantly enriched in genes functionally related to mitochondrial processes (Fig. 1d, Supplementary Table 4): *‘mitochondrial respiratory chain complex assembly* (FDR p-value < 1.21×10^-20^), *‘mitochondrion organization’* (FDR p-value < 9.05×10^-19^), *‘cytoplasmic translation’* (FDR p-value < 5.06×10^-17^), and *‘oxidative phosphorylation’* (FDR p-value < 6.60×10^-17^). Overall, our findings show that SIRT6 deficiency provokes significant gene expression changes in the mouse brain and induces transcriptional dysregulation of mitochondrial-related genes.

### SIRT6 regulates mitochondrial metabolism

The significant association of D.E. genes with essential mitochondrial processes allows us to speculate that SIRT6 silencing might also induce alterations in mitochondrial metabolite levels. To study the role of SIRT6 in mitochondrial metabolism, we examined the differential metabolite abundance patterns in SIRT6-KO untargeted LC-MS profiles compared to W.T. in the mouse embryonic stem cells (mESC) data (Supplementary Table 5). Similar to RNA-seq results, we observed a global difference between the abundance levels of W.T. and SIRT6-KO metabolite profiles, underlined by their clear separation by PC1 in the PCA plot (Fig. 2a). Differential abundance (DA) analysis revealed that 92 out of 235 metabolomic features (~39%, FDR < 0.05 and |log_2_ (Fold Change)|> 0.58) changed significantly between experimental conditions (Fig. 2b), including *Ascorbic acid* (upregulated), *Maleic acid* (downregulated), and *NAD^+^* (downregulated) as the most significant metabolites (Supplementary Fig. S2a,b). Consistent with the transcriptome analysis, we found a number of DA features related to mitochondrial energy system pathways. Fig. 2c, shows that several metabolites associated with catabolic process are more abundant in the SIRT6 W.T. group compared with SIRT6-KO, in four particular metabolites (*Malic acid, Fumaric acid, Oxoglutaric acid*, *Thiamine Pyrophosphate*) associated with TCA cycle and three metabolites (*NAD^+^*, *NADH, ADP)* associated with OXPHOS. The same tendency was observed for other DA metabolites related to the energy and carbohydrate metabolic pathways, of which only four metabolites were upregulated, while the rest fourteen were decreased in SIRT6-KO. In addition to these results, we found abundant alterations of metabolomic features that constitute the *Lipid* and *Amino Acid metabolism* pathways. Our results show that SIRT6 silencing alters cellular and mitochondrial metabolism.

**Figure 2:**
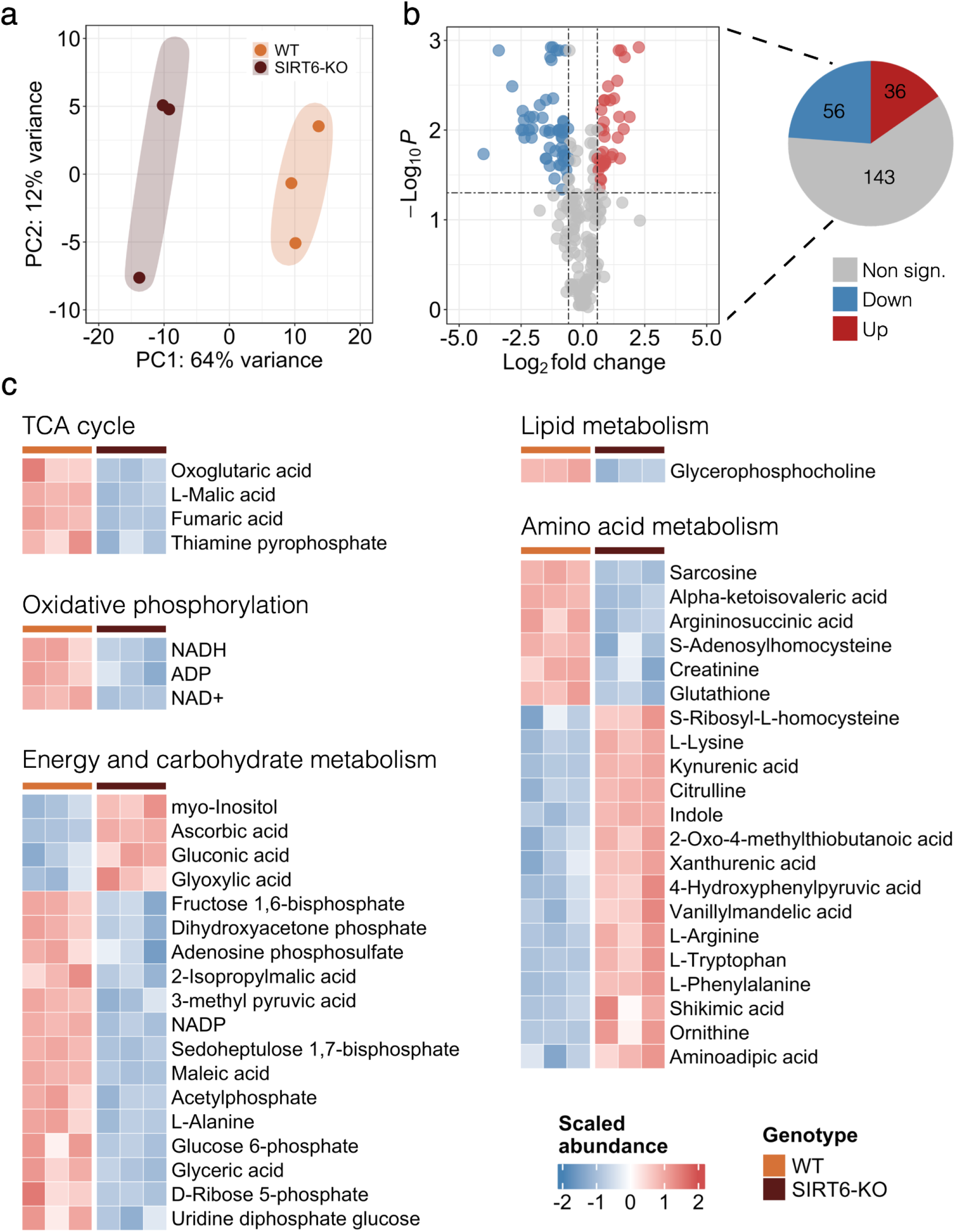
SIRT6 deficiency triggers an abundance of energy metabolites. **(a)** PCA plot showing separation between the W.T. (orange circles) and SIRT6-KO (brown circles) groups based on the mESC metabolomic profiles. Orange and brown ovals represent confidence ellipses of W.T. and SIRT6-KO groups. **(b)** The volcano plot illustrates differentially abundant metabolites detected between W.T. and SIRT6-KO mESC samples. Upregulated and downregulated metabolites are represented by red and blue circles, respectively. The pie plot (on the right) demonstrates the number of upregulated (red), downregulated (blue), and insignificant (gray) metabolites in the analysis. **(c)** The abundance heatmap of 68 out of 92 significant metabolites is classified according to the metabolic pathways they are involved in.

### SIRT6 deficiency leads to impaired oxidative phosphorylation

Furthermore, we focused on the significant D.E. mitochondrial genes that affected mitochondrial pathways. SIRT6 deficiency resulted in 256 significant D.E. mitochondria-related genes out of 1140 genes with confirmed mitochondrial localization according to the MitoCarta database^36^ (Fig. 3a). Importantly, downregulated genes constituted the majority (>91%) of all D.E. mitochondria-related genes. Of note, protein levels of one of the most significantly downregulated genes (*Cycs*, FDR p-value < 3.00×10^-19^) were consistently decreased in both male and female SIRT6-KO SH-SY5Y cells (Supplementary Figure 3a,b). On the contrary, both expression and protein levels of *Vdac1* changed insignificantly between female W.T. and SIRT6-KO samples but showed a reduction in protein levels in male SIRT6-deficient samples (Supplementary Figure 3a, b). Interestingly, we found that mitochondrial genes were overrepresented in their localization at the Mitochondrial Inner Membrane (MIM) compartment (Fig. 3b, Supplementary Fig. S32c). Given that MIM serves as a springboard for ATP synthesis, we hypothesize that significant mitochondria-related genes should be mostly associated with electron transport chain complexes. To explore biological functions related to D.E. mitochondrial genes, we performed G.O. enrichment analysis using information about mitochondrial pathways obtained from MitoCarta 3.0^36^ as a specific background. The top enriched categories included terms associated with mitochondrial respiratory chain complexes and mitochondrial ribosomes (Fig. 3c). Mitochondrial Complex I turned out to be the most affected by SIRT6 depletion (FDR p-value < 1.09×10^-07^), with 27 downregulated out of 43 genes encoding this Complex. Of note, 57 out of 99 genes encoding the electron transport chain subunits were differentially expressed in our analysis. Only *Succinate dehydrogenase complex flavoprotein subunit A* gene (*Sdha*) demonstrated an elevated level of expression (Fig. 3d). Also, we confirmed that these changes also occur in brain RNA-seq samples of two human donors from Allen Brain Atlas^37^, where the correlation between the expression of SIRT6 and the expression of OXPHOS-related genes is significantly stronger (p-value < 0.000636 and p-value < 0.000002, respectively) compared to other mitochondria-related genes (Supplementary Figure 3d).

**Figure 3:**
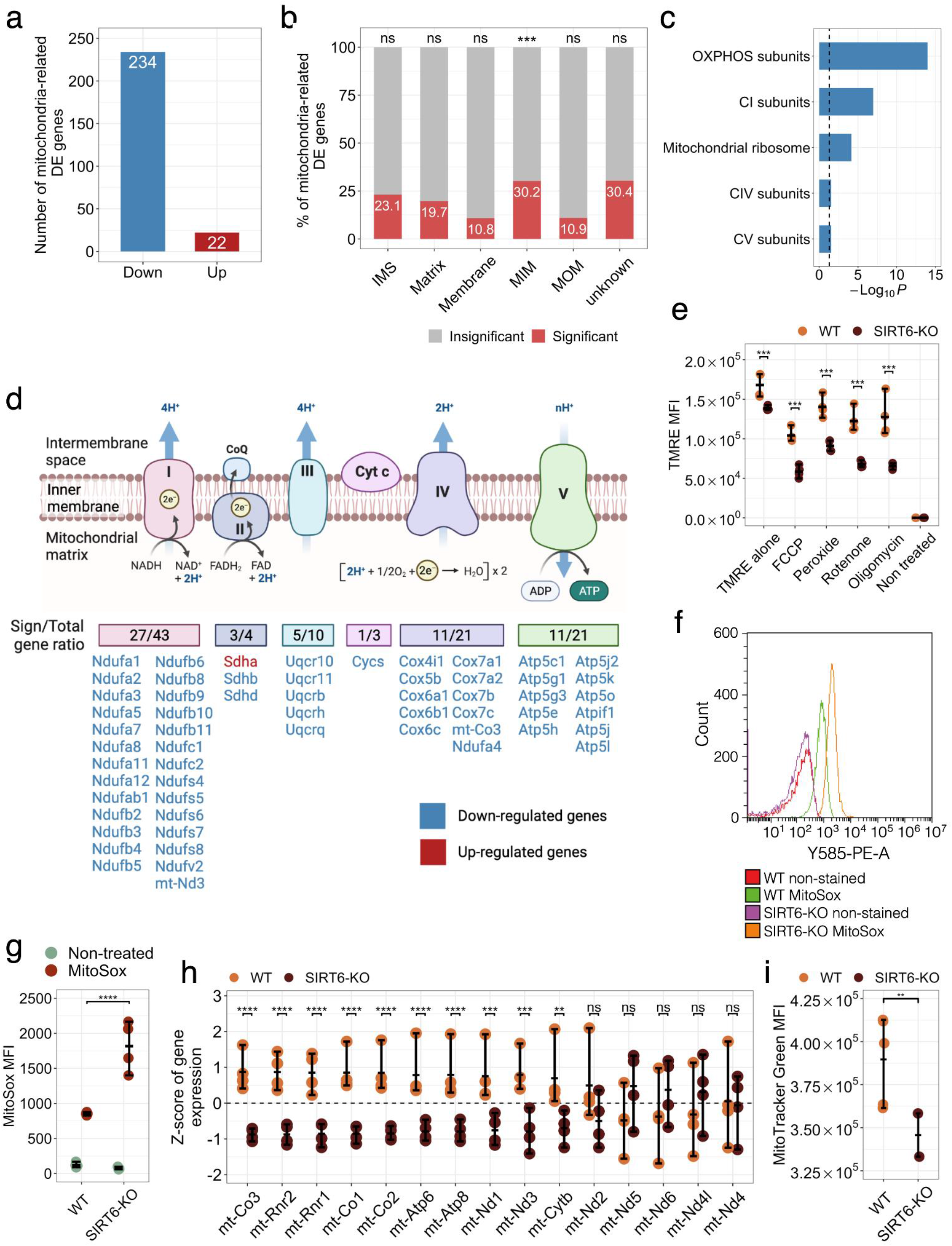
OXPHOS impairment in SIRT6-KO. **(a)** Number of significant D.E. genes associated with mitochondrial functions in W.T. compared to SIRT6-KO. Red and blue bars show the number of up-and downregulated genes. **(b)** Percentage of significant (red bars) and insignificant (gray bars) genes across mitochondrial compartments. ‘IMS’ denotes Intermembrane space, ‘MIM’ denotes mitochondrial inner membrane, and ‘MOM’ corresponds to the mitochondrial outer membrane. Black asterisks indicate the statistical significance of the enrichment (p-value<5.5×10^-04^, hypergeometric test). **(c)** Overrepresented mitochondrial pathways for W.T. compared to SIRT6-KO. The statistical significance threshold of 0.05 (hypergeometric test) is shown by a black dashed line. **(d)** Scheme illustrating the ratio between the number of significant D.E. genes associated with Cytochrome C oxidase and I-V complexes of the electron transport chain and the total number of genes per complex. Down and upregulated genes are marked by blue and red colors, respectively. **(e)** Mitochondrial membrane potential was measured in SH-SY5Y WT and SIRT6-KO cells under treatment with FCCP (10μM), Hydrogen peroxide (800 nM), Rotenone (200 nM), Oligomycin (20 μM), and without treatment. Asterisks indicate the level of statistical significance (p<0.05, t-test). **(f)** Mitochondrial ROS detection with MitoSox assay. The histogram shows fluorescence emission distributions measured in W.T., and SIRT6-KO cells that were non-stained and MitoSox treated. Distribution of mean fluorescence intensity (MFI) values measured in W.T., and SIRT6-KO cells that were non-treated (green circles) and MitoSox treated (red circles). Error bars are mean ± SD **** p<0.0001 (n=3-5). **(h)** Z-score transformed expression levels of mtDNA genes detected in our RNA-seq experiment. Orange and brown circles represent W.T. and SIRT6-KO samples, respectively. **(i)** Difference in mitochondrial content between W.T. and SIRT6-KO SH-SY5Y cells. Asterisks indicate the level of statistical significance (p<0.05, t-test).

We hypothesized that a global reduction in the expression of OXPHOS genes and electron transport chain (ETC) complex activity in SIRT6-KO models might be accompanied by the corresponding decline in mitochondrial membrane potential ΔΨ. In order to check this hypothesis, we first measured ΔΨ in W.T. and SIRT6-deficient SH-SY5Y cells stained with TMRE dye. Indeed, SIRT6-KO mitochondria revealed a significant 1.21-fold decrease in ΔΨ compared to W.T. cells (FDR p-value < 0.0006, Tukey’s multiple comparisons test, Fig. 3e, Supplementary Table 6). Then, we tested ΔΨ in the same W.T. and SIRT6-KO cells but treated with an uncoupler of oxidative phosphorylation FCCP. Interestingly, supplementation of FCCP enhanced the reduction effect of ΔΨ upon SIRT6 deficiency, resulting in 1.78 fold decrease of ΔΨ in SIRT6-KO cells (FDR p-value < 0.0001, Tukey’s multiple comparisons test). Similar significant changes were observed when inhibitors of individual complexes of the ETC were added to the cells. SIRT6-KO cells with inactivated Cytochrome C complex by hydrogen peroxide showed 1.54 fold reduction in ΔΨ (FDR p-value < 0.0001, Tukey’s multiple comparisons test), while mitochondria with inactivated Complex I (rotenone treatment) and ATP synthase (oligomycin treatment) showed the highest level of ΔΨ reduction in SIRT6-KO, in 1.81 and 1.93 times, respectively (FDR p-value < 0.0001 and FDR p-value < 0.0001, Tukey’s multiple comparisons test), suggesting higher dependence of SIRT6 for these complexes. Then, we speculated that an elevated ROS production could also accompany observed transcriptional changes of OXPHOS-related genes and ΔΨ reduction upon SIRT6 knockout. Indeed, using MitoSox staining, we detected significantly high levels of ROS in SIRT6-KO cells compared to W.T. (Fig 3f,g). All these results collectively indicate that the mitochondrial oxidative phosphorylation process is markedly impaired in SIRT6-deficient cells.

### Lack of SIRT6 results in a reduction of mtDNA gene expression and mitochondrial content

Mitochondrial activity is regulated by both nuclear and mitochondrial DNA encoded genes. Since all mitochondrial-encoded genes are involved in oxidative phosphorylation, we studied the expression changes of these genes in SIRT6-KO brains. In particular, we extracted the expression of 15 mtDNA genes detected in our RNA-seq data and the direction of their expression changes in W.T. and SIRT6-KO samples. Four out of these 15 mtDNA genes were downregulated in SIRT6-KO mice, including statistically significant genes *mt-Co3* (FDR p-value < 3.8×10^-18^), *mt-Rnr2* (FDR p-value < 1.1×10^-14^), *mt-Rnr1* (FDR p-value < 1.0×10^-11^), *mt-Nd3* (FDR p-value < 5.2×10^-04^) (Fig. 3h). In addition, six other mtDNA-encoded genes (*mt-Co1, mt-Co2, mt-Atp6, mt-Atp8, mt-Nd1, mt-Cytb)* showed a statistically significant reduction in expression (FDR < 0.05), but did not meet log_2_ (Fold Change) criterion for significance. Since altered mtDNA gene expression levels might indicate co-directional changes in mitochondrial content, we also measured mitochondrial mass in W.T. and SIRT6-KO SH-SY5Y cells using the MitoTracker Green assay. Consistent with transcriptional down-regulation patterns of mtDNA genes, mitochondrial mass was significantly lower in SIRT6-deficient cells (~21.8% decrease, T-test p-value < 0.0087) than in W.T. cells (Fig. 3i, Supplementary Table 6), which in turn can be a marker of impaired mitochondrial biogenesis or increased degradation.

### SIRT6-SIRT4 and SIRT6-YY1 axes promote OXPHOS in the brain

Next, we elucidated the mechanism behind the SIRT6-dependent regulation of mitochondrial activity and the oxidative phosphorylation process. First, we explored SIRT3, SIRT4, and SIRT5 genes from the sirtuin family, which encode proteins localized in mitochondria and coordinately impact mitochondrial pathways related to redox homeostasis and cellular metabolism^38^. To determine whether SIRT6 may transcriptionally regulate these genes, we examined their expression patterns upon SIRT6 knockout (Fig. 4a). SIRT3 and SIRT4 were significantly reduced in SIRT6-KO brains (FDR p-value < 3.60×10^-12^ and 3.33×10^-06^, respectively). At the same time, the lack of SIRT6 did not substantially affect SIRT5 expression. We further confirmed the positive association between SIRT6 and SIRT3-4 by analyzing publicly available gene expression data in the mouse brain from Zhang et al.^39^ (Fig. 4b). SIRT6 expression levels positively correlated with the corresponding expression levels of all mitochondrial sirtuins (Pearson’s R = 0.5, 0.79, 0.71 for correlations with SIRT3, SIRT4, SIRT5, respectively). Then, we focused on SIRT3 and SIRT4 genes, which most significantly changed among mitochondrial sirtuins. To experimentally validate their role in OXPHOS regulation, we assessed the changes in mitochondrial membrane potential ΔΨ between W.T. and SIRT6-KO SH-SY5Y cells with overexpressed SIRT3 and SIRT4. We found that increased expression of SIRT3 partially rescued ΔΨ in SIRT6-deficient cells (Fig. 4c). On the contrary, ΔΨ in SIRT6-KO cells coupled with SIRT4 overexpression was significantly recovered compared to that in W.T. cells, suggesting that SIRT4 is of greater importance for the regulation of oxidative phosphorylation when SIRT6 is absent.

**Figure 4:**
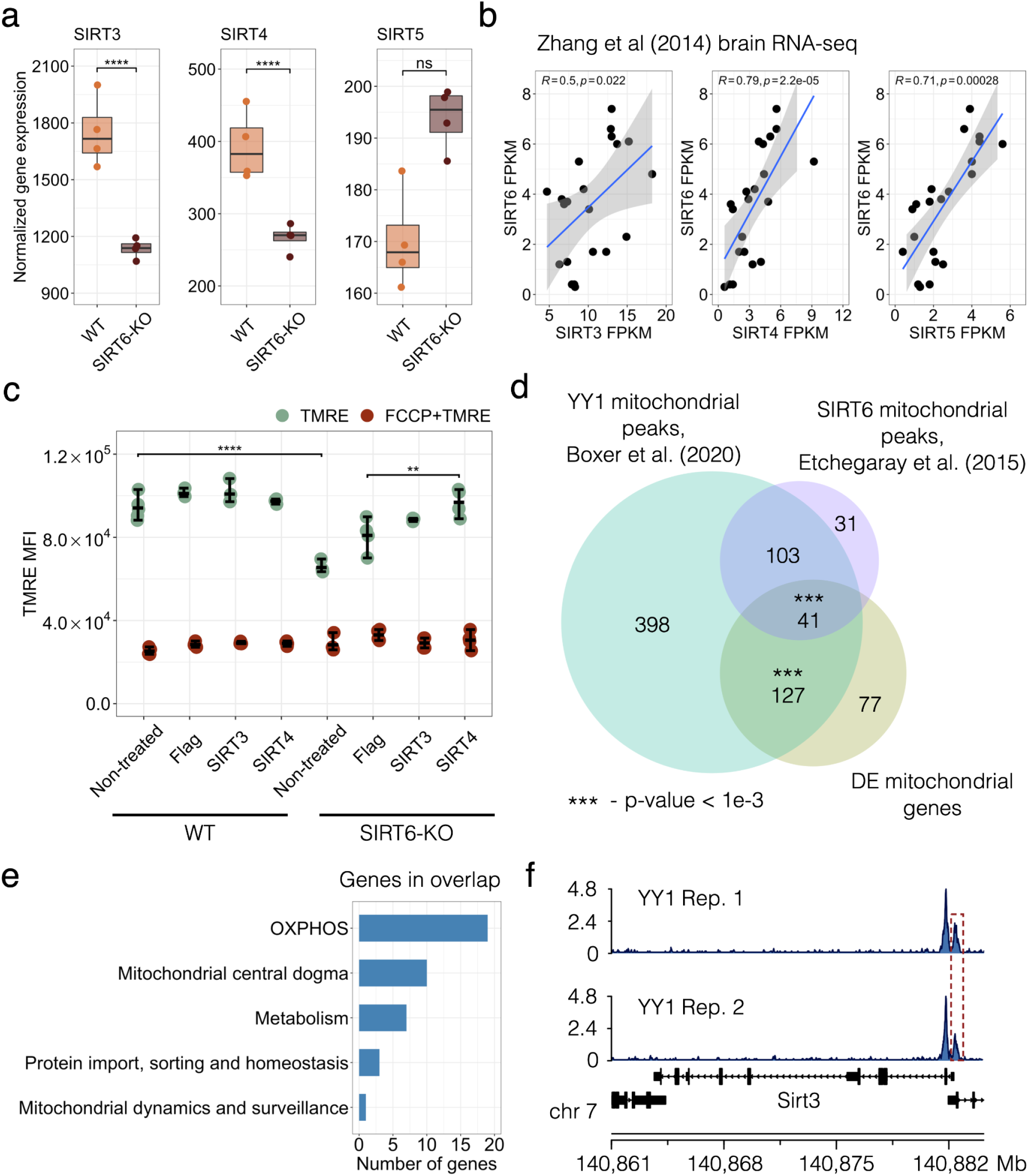
SIRT6-SIRT4 and SIRT6-YY1 axes in mitochondrial regulation. **(a)** SIRT3-5 expression levels in transcriptomic profiles of W.T. and SIRT6-KO mice. Asterisks indicate the statistical significance (p-value < 0.05) derived from DESeq2 differential expression analysis. **(b)** Spearman’s correlations of SIRT6 expression profile with expression profiles of mitochondrial sirtuins (SIRT3-5) in the brain RNA-seq data from Zhang et al.^39^. **(c)** Mitochondrial membrane potential measured in SH-SY5Y WT and SIRT6-KO cells when SIRT3 or 4 were exogenously overexpressed. W.T. and SIRT6-KO SH-SY5Y cells were transfected with Flag-CMV, SIRT3-Flag-CMV, and SIRT4-Flag-CMV plasmids. After 48 hours, cells were collected and stained with TMRE and Life/dead viability dye and the intensity of fluorescence was measured by CytoFLEX. **(d)** Venn diagram showing overlaps between significant mitochondria-related genes from the RNA-seq analysis (orange), YY1 mitochondrial targets (green) and SIRT6 mitochondrial targets (purple). Statistical significance of the overlaps was calculated using the permutation test. **(e)** Bar plot showing biological functions along with the number of the mitochondria-related genes overlapped between all datasets presented in Fig. 4d. **(f)** YY1 peaks at SIRT3 promoter in two analyzed ChIP-seq replicates.

Another gene we explored in the context of the OXPHOS regulation was YY1. We have previously shown that SIRT6 and YY1 can act together, forming a complex that regulates several shared target genes^24^. To examine whether they can also regulate mitochondrial processes in a coordinated manner, we analyzed two publicly available YY1 ChIP-seq in cortical neurons (GSE128182 GEO accession). We searched for YY1 peaks corresponding to the promoters of mitochondria-related genes. In addition, we compared these peaks with both mitochondria-related D.E. genes from our RNA-seq analysis as well as with SIRT6 targets involved in mitochondrial regulation derived from public mESC ChIP-seq profiles (GSE65836). As a result, we detected 669 YY1 peaks associated with promoters of mitochondrial genes, including 168 peaks that were localized within 1 kb from the promoters of mitochondrial D.E. genes and 144 peaks colocalized with SIRT6 binding sites in mESC (Fig. 4d). We also identified only 11 SIRT6 binding sites in the absence of YY1 peaks at mitochondria-related gene promoters, also suggesting a smaller indirect mechanism of mitochondrial regulation by SIRT6.

Interestingly, both YY1 and SIRT6 peaks were overrepresented at the promoters of genes localized in the mitochondrial protein-containing complex (FDR p-value < 1.01×10-38 and FDR p-value < 3.96×10-12, respectively) and the mitochondrial inner membrane (FDR p-value < 1.82×10-27 and FDR p-value < 2.66×10-08, respectively), while YY1 target genes were also enriched for mitochondrial matrix (FDR p-value < 3.41×10-37) and ATPase complex (FDR p-value < 4.34×10-22) (Supplementary Fig. S4a, b). Our analysis revealed that the expression of more than 66% of the detected mitochondria-related genes could be regulated by either YY1 or by YY1 and SIRT6 together. Both YY1 and SIRT6 peaks were found within promoters of 41 mitochondria-related D.E. genes that were also over-represented (permutation test p-value < 5.1×10^-04^) in this overlap compared to non-significant mitochondria-related genes. These genes are also related to OXPHOS, mitochondrial metabolism, and protein import regulation (Fig. 4e, Supplementary Fig. S4c). Besides its coordinated regulatory activity with SIRT6, YY1 can also independently bind to the promoters of mitochondria-related D.E. genes. In our analysis, it was enriched (permutation test p-value < 1.0×10-^03^) at the promoters of such 127 D.E. genes, including SIRT3 (Fig. 4f), whose importance for OXPHOS was shown above. Hence, our analysis suggests that SIRT6 may act as a regulator of mitochondrial functions via the SIRT6-YY1-SIRT4 axis.

### Neuropathological role of SIRT6 through the prism of mitochondrial deregulation

Sirtuin 6 has been reported to be important in the protection against age-related and neurodegenerative diseases in the brain^18,23,40,41^. Since, in our analysis, we observed a global reduction in the transcriptional level of mitochondrial genes, we explored whether these changes can be linked to pathways of age-associated diseases occurring in the brain. Therefore, we performed the Gene Set Enrichment Analysis (GSEA) based on all genes in our RNA-seq dataset. This analysis revealed 71 significantly affected KEGG pathways (Supplementary Table 7, Supplementary Fig. S5), including ‘Parkinson’s disease (FDR p-value < 0.015), ‘Huntington’s disease.

(FDR p-value < 0.0168), ‘Alzheimer’s disease (FDR p-value < 0.0169), and ‘Amyotrophic lateral sclerosis (FDR p-value < 0.0168) pathways (Fig. S5). Interestingly, these neurodegenerative disease pathways formed one distinct cluster with ‘Oxidative phosphorylation’ pathway in the enrichment network, showing a large number of overlapping genes between them (Fig. 5a).

**Figure 5:**
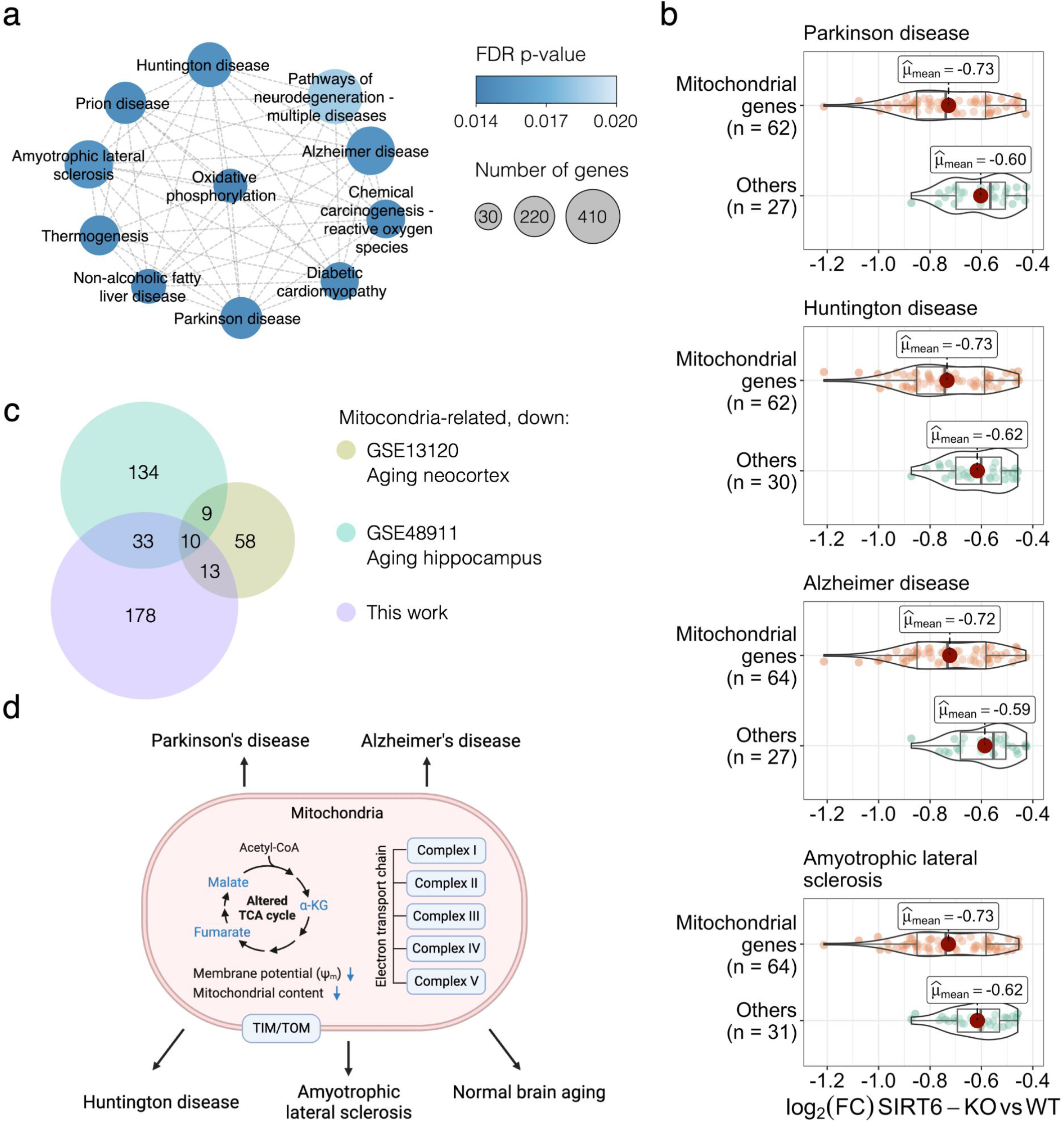
SIRT6 silencing triggers neurodegenerative disease pathways and normal brain aging. **(a)** Cluster of enriched KEGG pathways obtained using GSEA. Each circle represents an enriched pathway and is colored according to the FDR p-value. **(b)** Violin plots representing the log_2_ (Fold Change) expression for genes with the largest contribution to the GSEA enrichment result per neurodegenerative pathway. **(c)** Euler diagram showing ten common downregulated genes between publicly available aging brain gene expression datasets (GSE13120 and GSE48911 accessions in the GEO database) and significantly downregulated mitochondrial genes from this study. **(d)** Proposed model of the mitochondrial dysfunction caused by SIRT6 silencing and its involvement in neurodegenerative diseases and normal aging. The figure was generated using the BioRender website [https://app.biorender.com/biorender-templates].

To check whether the expression changes of mitochondria-related transcripts directly caused the enrichment of these pathways, we retrieved core enrichment genes from the paths of interest. More than 67% of core enrichment genes in each selected pathway were associated with mitochondrial functions. The highest percentage was detected for Alzheimer’s disease (Fig. 5b). Moreover, the mitochondria-related core enrichment genes exhibited lower mean log_2_ (Fold Change) values compared to non-mitochondrial genes in each of the selected neurodegenerative disease pathways.

Since mitochondrial dysfunction is one of the most stable and crucial hallmarks of normal aging^25,26,42^, we then compared our downregulated mitochondria-related genes with genes that were previously reported to be signatures of mouse brain aging^43,44^. As a result, we captured ten downregulated mitochondrial genes that also showed a reduction in their expression levels in both neocortex and hippocampus aging data (Fig 5c). Notably, this list of commonly downregulated genes included genes related to OXPHOS complexes (*Sdhd, Ndufa7, Uqcrq)* and mitochondrial protein import machinery (*Timm10b)*. Another interesting overlapping gene was *Uracil DNA Glycosylase (Ung)*, which has an important role in mitochondrial base excision repair (BER) initiation. Given the limited DNA repair mechanisms in mitochondria, one can expect that the decrease in *Ung* activity might provoke an accumulation of mutations in mtDNA. However, this question requires further investigation. Taken together, the findings reported above suggest that the brain lacking SIRT6 expression is characterized by mitochondrial dysfunction (OXPHOS impairment, TCA dysregulation, reduced mitochondrial content and membrane potential) that causes a neurodegenerative-like phenotype and contributes to a pathological aging of the brain.

## Discussion

In this study, we performed RNA-seq, GC-MS, and LC-MS experiments to trace molecular organization changes at transcriptome and metabolome levels of SIRT6 knockout systems. We show that SIRT6 deficiency leads to a dramatic downregulation of mitochondrial genes and changes in mitochondria-related metabolite composition, suggesting that SIRT6 critically regulates mitochondrial activity in the brain.

Our analysis of gene expression levels in the SIRT6-deficient mouse brain revealed dramatic transformations upon SIRT6 knockout: almost three thousand genes changed their expression significantly, with a strong enrichment of mitochondrial functions among down-regulated ones, allowing us to conclude that SIRT6 knockout induces transcriptional dysregulation of mitochondrial genes. This result bridges the missing gap between studies demonstrating mitochondrial dysfunction in normal and pathological aging ^1,45^ and studies proclaiming the critical role of SIRT6 in the protection against aging-associated diseases^18,19^.

Though mitochondrial dysfunction is a known marker of aging and neurodegenerative diseases, the exact mechanism behind it remains unknown. Our study suggests that the decay of SIRT6 levels during aging^18^ and in Alzheimer’s disease^18,23,46^ could be a key motive in the deterioration of mitochondrial functions. The changes induced by the SIRT6 knockout that we observe at the metabolite level support this claim: metabolites related to mitochondrial energy system pathways (in particular, OXPHOS and TCA cycle) are significantly overrepresented among differentially abundant metabolites. In line with the discussed mitochondrial dysfunction in aging, all but one of these metabolites are downregulated in the SIRT6-KO samples. Importantly, the dramatic decline of one of them, NAD+, was also associated with pro-senescence mechanisms in various species^47,48^, as well as with limited neuroprotective activity of sirtuins^49^.

Accordingly, the vast majority of differentially expressed mitochondria-related genes were downregulated in our gene expression analysis. As they were strongly enriched with mitochondrial respiratory chain complexes, we measured the mitochondrial membrane potential and mitochondrial content in SIRT6-KO cells because reduced gene expression might indicate the loss of mitochondria. Both measured characteristics were significantly decreased, validating the suggested impairment of mitochondrial oxidative phosphorylation and mitochondrial biogenesis in SIRT6-deficient brains. Interestingly, the average decrease of mtDNA gene expression (~19.7%) in SIRT6-KO was in good agreement with the corresponding reduction of mitochondrial content (21.8%), suggesting impaired mitochondrial biogenesis as a primary cause of the observed transcriptional dysregulation in mitochondria upon SIRT6 knockout.

Concordantly, the impaired membrane potential upon SIRT6-KO can be partially rescued by restoring SIRT3 and SIRT4 levels, which were significantly downregulated in SIRT6-deficient brains. Both of them appear to be a target of SIRT6, localized in mitochondria and impact on mitochondrial pathways related to redox homeostasis and cellular metabolism^38^. The analysis of our and publicly available gene expression data^39^ confirms that SIRT6 transcriptionally regulates SIRT3 and SIRT4. Both of which have important roles in mitochondria metabolism ROS balance and lifespan^50–52^. Another companion of SIRT6 is the transcription factor YY1. We have shown them to act together, forming a complex that regulate many shared target genes^24^. Our analysis of YY1 ChIP-seq data^53^ suggests that SIRT6 and YY1 regulate mitochondrial processes coordinately.

Taken together, our results show that SIRT6 knockout induces global molecular transformations in the brain: almost three thousand genes change their expression significantly, as well as nearly half of all studied metabolites. Part of these differences are distinctly attributed to mitochondrial dysfunction, particularly in mitochondrial respiratory chain complexes, as confirmed by measurements of the mitochondrial membrane potential and mitochondrial content. This SIRT6-dependent regulation of mitochondrial activity might be coordinated with SIRT3 and YY1. Lastly, we confirmed that these changes occurred in neurodegenerative diseases and aging brains, pointing to SIRT6 decay as the main reason for these changes.

## Materials and Methods

### Generation of brain-specific SIRT6-KO mice and cells

SIRT6-KO mice and SH-SY5Y SIRT6-KO cells were generated using the protocol described in Sebastián et al.^54^ and Kaluski et al.^18^.

### RNA preparation and quality control

RNA was extracted from mice’s left brain hemispheres, using the NucleoSpin RNA Plus kit (MACHEREY-NAGEL GmbH & Co. K.G., catalog number 740984.50), according to the manufacturer’s manual. The purified RNA was then cleaned from any possible residual genomic DNA contamination using the RNeasy MinElute Cleanup Kit (QIAGEN, catalog number 74204), according to the manufacturer’s manual. Using TapeStation, RNA Integrity Number (RIN) was then assessed and only samples with RIN>8.7 were in use.

### Full-length poly-A RNA sequencing

Library preparation was conducted by The Crown Genomics Institute of the Nancy and Stephen Grand Israel National Center for Personalized Medicine, Weizmann Institute of Science, Israel (G-INCPM). Briefly, the library kit used was the in-house INCPM mRNA-seq kit (G-INCPM, Weizmann Institute of Science), for full-length RNA-seq with polyA-based capturing. Sequencing was done using 2 lanes of NextSeq 500 High Output v2.5 Kit (75 cycles) (Illumina Inc., catalog number 20024906).

### RNA-seq data processing

Raw reads from 8 *M. musculus* RNA samples were filtered and trimmed using *fastp*^55^ and then processed via version 3.0 of the *nf-core/rnaseq* pipeline [DOI: 10.5281/zenodo.1400710]. In brief, trimmed reads were filtered with *Trim Galore!* tool and mapped to the mouse GRCm39 reference genome with *STAR*^56^. Then gene expression was quantified using the Salmon tool^57^. The full guidelines for the pipeline are available at https://nf-co.re/rnaseq. Gene annotation was performed using the *AnnotationDbi* R package [DOI: 10.18129/B9.bioc.AnnotationDbi] with downloaded *EnsDb.Mmusculus.v79* annotation database [DOI: 10.18129/B9.bioc.EnsDb.Mmusculus.v79] generated from Ensembl.

### Differential expression analysis

Differential expression (D.E.) analysis was performed via the DESeq2^58^ package in the R programming language. First, we removed low-expressed genes for which the minimum expression level within any group of samples was < 3. Then raw gene counts were normalized using DESeq2’s median of ratios method and quality control procedures were performed. The following design formula was used to evaluate expression differences between groups of samples: *design = ~ genotype*. After fitting the Negative Binomial model for each gene, we performed pairwise comparisons between groups using the Wald test. Genes were considered to be differentially expressed if *FDR p-value* < 0.05 and *|log2 (Fold Change)|* > 1.5.

### Functional analysis of genes

We used *clusterProfiler* R package^59^ to perform Gene Ontology (G.O.) enrichment analysis on both sets of down- and upregulated genes. Redundant G.O. categories were removed using the ‘simplify’ function from *clusterProfiler* package with default settings. Gene Set Enrichment Analysis (GSEA) was performed with *gseaKEGG* function from *clusterProfiler* and pairwise similarity of significant KEGG pathways was calculated with *pairwise_termsim* function from *enrichplot* package [DOI: 10.18129/B9.bioc.enrichplot] using Jaccard similarity measure. Then the pathway similarity network was constructed and visualized in *Cytoscape*^60^. Next, core enrichment genes for four pathways related to neurodegenerative diseases (Parkinson’s disease, Huntington’s disease, Alzheimer’s disease, Amyotrophic lateral sclerosis) were retrieved and classified in two groups according their relevance to mitochondria (‘mitochondrial genes’ and ‘others’ groups). Expression level distributions for these two groups were visualized using *ggstatsplot* R package [DOI: https://doi.org/10.21105/joss.03167].

### Analysis of mitochondria-related genes

A list of mouse mitochondria-related genes, as well as information regarding their sub-mitochondrial localization and related mitochondrial pathways, were obtained from the *MitoCarta* database (version 3.0) ^36^. A total of 149 mitochondrial pathways were used for the enrichment analysis of D.E. mitochondria-related genes, performed with the ‘enricher’ function from the *ClusterProfiler* package. An illustration of electron transport chain complexes with associated D.E. genes was performed using the *BioRender* website [https://biorender.com/].

### Extraction, measurement of metabolomics profiles and data processing

Extraction and measurement of polar metabolites from brain tissue using LC-MS and GC-MS was performed as described by Giavalisco P. et al (2011) and Lapidot-Cohen et al. (2020). In brief, to approximately of brain tissue, 1 ml of a homogeneous mixture of pre-cooled methanol/methyl-tert-butyl-ether/water (1:3:1) were added and votexed. This was followed shaking for 10 min and another 10 min of incubation in an ice-cooled ultrasonication bath. 500 μl of UPLC-grade methanol/water (1:3) was added to homogenate, then vortexed and spun for 5 min at 4°C. The result was a phase separation with polar and semi-polar metabolites in the lower aqueous phase. Equal volume of that phase 300 μl was taken twice: to two separate tubes and next dried down in Speedvac and stored at −80°C until subsequent L.C.–M.S. and GC-MS analysis. Prior to LC-MS analysis, samples were re-suspended in 80% (v/v) methanol and 20% (v/v) water. L.C.–M.S. data were obtained using a Waters Acquity UPLC system (Waters), coupled to an Exactive mass spectrometer (Thermo Fisher Scientific). A HSS T3 C18 reversed-phase column (100mm×2.1mm×1.8μm particles; Waters) was used and the temperature was set to 40°C. The mobile phases were 0.1% formic acid in H_2_O (Buffer A, ULC MS grade; BioSolve) and 0.1% formic acid in acetonitrile (Buffer B, ULC MS grade; BioSolve). A 5-μl sample was injected. The spectra were recorded alternating between full-scan and all-ion fragmentation-scan modes, covering a mass range from 100 to 1500 *m/z*. The resolution was set to 25000, with maximum time scan 250 ms. Chromatograms from the UPLC-FT/MS runs were analyzed and processed with Compound Discoverer 3.3 (Thermo Fisher Scientific) and Xcalibur^™^ Software (Thermo Fisher Scientific). Molecular masses, retention times, and associated peak intensities for each sample were extracted from the .raw files. Prior to GC-MS analysis samples were derivatized (Lisec et al., 2006) and metabolites were measured by G.S.−TOF–M.S. (Leco Pegasus HT TOF-MS). Gas chromatography was performed on 30 m DB-35 column with helium as a carrier gas with a constant flow rate of 2 ml s^-1^. Injection temperature was 230°C while the transfer line and ion source were set to 250°C. 85°C was the initial temperature of the oven, increased at a rate of 15°C min^-1^ up to a final temperature of 360°C. Mass spectra were recorded at 20 scans s^-1^ with m/z 70-600 scanning range. Retention times, molecular masses and associated peak intensities for each sample were extracted from the .cdf files. Targeted analysis was performed by XcaliburTM Software (Thermo Fisher Scientific) and supported by TargetSearch.

### Differential abundance analysis

Metabolite differential abundance analysis was done with the MetaboAnalyst platform^61^. Annotated mouse ESC metabolites were normalized via the median normalization method and then were log_2_ transformed. Principal components of the data were calculated using the ‘prcomp’ function in R and then used for the visualization of the profiles. Student’s t-test was applied to define significantly changed metabolites, followed by the log_2_ (*Fold Change*) calculation. Differentially accumulated metabolites (DAMs) were retrieved according to *FDR < 0.05* and |log_2_ (*Fold Change)|* > 0.58 cutoff criteria. Volcano plot visualization was done with the EnhancedVolcano [DOI: 10.18129/B9.bioc.EnhancedVolcano] package in R. Significant features were classified by their metabolic pathway identity provided by the KEGG database^62^. ComplexHeatmap R package^63^ was used to plot heatmaps of metabolite abundances.

### Analysis of public brain RNA-seq data

Processed and FPKM normalized mouse brain RNA-seq profiles were downloaded from Zhang et al.^39^. Then only expression levels of SIRT6 and mitochondrial sirtuins (SIRT3, SIRT4, SIRT5) were selected, followed by Spearman’s correlation calculation. Analysis of correlation of SIRT6 with mitochondria-related genes was done using brain RNA-seq data of two human donors (H0351.2001, H0351.2002) from Allen Brain Atlas database^37^. Using the list of mitochondria-related genes retrieved from MitoCarta (version 3.0), Spearman’s correlations of SIRT6 with OXPHOS and non-OXPHOS related genes were calculated for both donor expression profiles. Permutation test (n permutation = 1000000) was used to test the assumption regarding unlikeness of distributions for OXPHOS and non-OXPHOS related genes.

### Analysis of YY1 and SIRT6 ChIP-seq data

Processed data of two mouse YY1 ChIP-seq replicates in cortical neurons (SRX5509061 and SRX5509062 accession numbers^53^, and SIRT6 ChIP-seq replicates in mouse embryonic stem cells (SRX873340, SRX873342, SRX873343 accession numbers^64^) were downloaded from the *ChIP-Atlas* database ^65^. Called peaks with q < 1×10^-05^ were annotated by their genome position using ‘annotatePeak’ function from *ChIPseeker* package^66^ and only peaks localized at promoters of mitochondria-related genes in all replicates were selected. SIRT6 peaks called for both SIRT6-KO and W.T. replicates were subtracted from the analysis. The cellular component (CC) G.O. analysis of genes associated with the selected YY1 and SIRT6 peaks was performed using ClusterProfiler. Area-proportional Venn diagram for mitochondria-related D.E. genes, YY1 and SIRT6-regulated mitochondria-related genes was plotted using *venneuler* R package [https://cran.r-project.org/package=venneuler]. The significance of overlap was calculated via a permutation test with non-significant mitochondria-related genes as a specific background. ChIP-seq profiles of the selected peaks were visualized with the *karyoploteR* package^67^.

### Analysis of public aging brain datasets

Microarray gene expression data of mouse aging neocortex (five 5-month-old and five 30-month-old, GSE13120^43^) and mouse aging hippocampus (three 10 days old and three 20-month-old, GSE48911 ^44^) were used for the analysis. Only WT replicates were selected from aging hippocampus datasets. Differential expression analysis was performed via *GEO2R* online tool^68^ with default parameters.

### Quantification of mitochondrial mass

To measure the mitochondrial mass, SH-SY5Y SIRT6-Control and SIRT6-KO cells were stained 100 nM MitoTracker Green (M7514; Molecular Probes) and viability dye (eBioscience^™^ Fixable Viability Dye eFluor^™^ 780, 65-0865-14, Invitrogen) in the dark for 30 min at 37°C. Fluorescence intensities of Mitotracker^™^ Green were detected by using flow cytometr (CytoFLEX S Flow Cytometer).

### Mitochondrial Membrane Potential Assay

To measure the mitochondrial membrane potential, SH-SY5Y SIRT6-Control and SIRT6-KO cells were treated with trypsin, washed once and 1×10^5^ cells were loaded to each well of 96 well plate. Cells were resuspended in DMEM with 10% FBS containing different concentrations of rotenone or oligomycin or hydrogen peroxide or FCCP as indicated in figure legends and incubated in the dark for 30 min in cell culture incubator. After 30 min, cells were washed in PBS and incubated in the dark for 30 min with TMRE and viability dye at 37°C (TMRE Mitochondrial Membrane Potential Assay Kit, 701310). After cells were washed and fluorescence intensities were detected by using flow cytometry (CytoFLEX S Flow Cytometer).

### Measurement of mitochondrial ROS generation

The measurement of mitochondrial ROS generation was performed by using MitoSOX^™^ Red staining. SH-SY5Y SIRT6-Control and SIRT6-KO cells were treated with trypsin and then centrifuged at 500 g for 5 min. Cells were washed once with HBSS and incubated with 5 μM MitoSOX^™^ and viability dye for 15 min at 37 °C in dark. After treatment cells were washed in HBSS and MitoSOX^™^-positive cells were detected by using flow cytometry (CytoFLEX S Flow Cytometer).

### Western blot analysis

Total protein extracts from WT (n=8) and brSirt6KO (n=8) brains were prepared. Fifteen micrograms of protein were loaded onto 4-20% Tris-Glycine polyacrylamide gel (BioRad). Proteins were separated for 1 hr at 120 V and then blotted to nitrocellulose membranes at 100 V for 35 min. The blots were blocked with 5% BSA in TBST (15 mM Tris-HCl, pH 7.5, 200 mM NaCl, and 0.1% Tween 20) for 1 hr at room temperature. Membranes were incubated ON with a mouse monoclonal anti-cytochrome c antibody (clone 7H8.2C12; 1:1000; PharMingen Becton Dickinson), a mouse monoclonal anti-VDAC1 antibody (Santa Cruz biotechnology, sc-390996) and anti-β-tubulin antibody (Cell Signaling, 2128). The blots were developed using the Western Bright Quantum HRP substrate chemiluminescence reagent (K-12042, Advansta).

## Data availability

Raw RNA-seq data described in the study is uploaded to the Gene Expression Omnibus (GEO) database under X accession.

## Acknowledgments

RNA-seq data processing was performed on the Arkuda HPC cluster provided by Skolkovo Institute of Science and Technology.

## Funding

The study was funded by the David and Inez Myers foundation, the Israeli Ministry of Science and Technology (MOST), the High-tech, Bio-tech and Negev fellowships of Kreitman School of Advanced Research of Ben Gurion University, and the RNA-seq analysis was supported by the Russian Science Foundation (grant number 21-74-10102 to E.K.).

## Author contributions

DiS performed all the bioinformatic analysis and wrote the paper EE performed molecular biology experiments, DaS and SK performed animal experiments and RNA-seq WJ and YB performed metabolomic experiments in SHSY5Y with SK, CC, BMP and RM performed metabolomic assay on ES cells EK guided bioinformatic analysis and wrote the paper DT planed the project advise on experiments and results and wrote the paper.

## Competing interest statement

The authors declare no competing financial interests.

## Supplemental information

**Supplementary figure 1:**
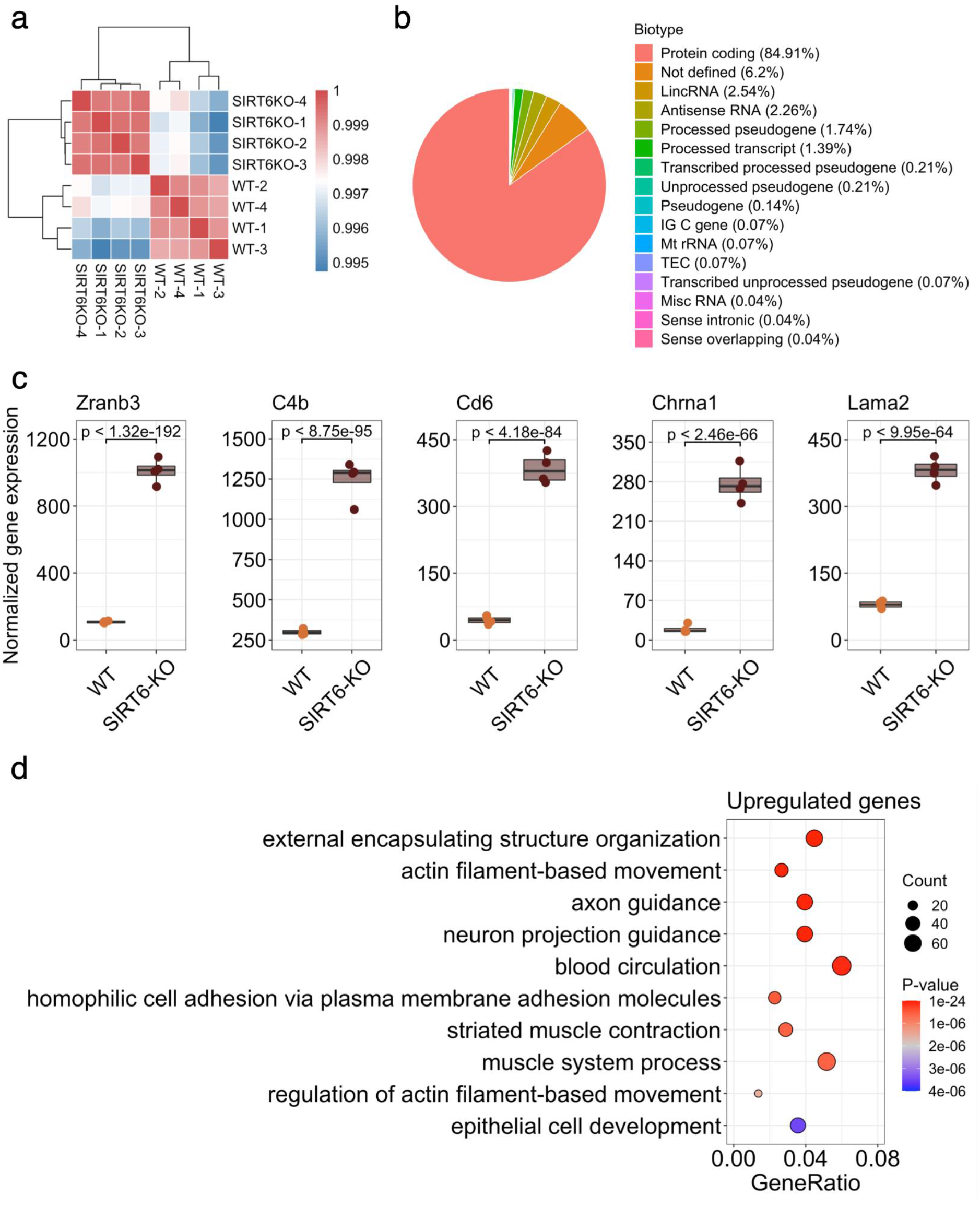
Analysis of WT and brSIRT6-KO gene expression data (related to Fig. 1). **(a)** Heatmap illustrating Pearson’s correlation between experimental samples. **(b)** Pie plot representing proportion of biotype annotations among DE genes. ‘IG C gene’ annotation denotes constant chain immunoglobulin genes, ‘TEC’ annotation describes predicted genes that require experimental validation. **(c)** Boxplots of the top 5 most differentially expressed genes in the analysis. Expression of WT samples are shown by orange points and expression of brSIRT6 are shown by brown points. **(d)** GO analysis showing top 10 enriched biological processes for upregulated genes. Each circle corresponds to the enriched GO term and varies in size according to the number of significant genes belonging to this term. Gene ratio represents the number of DE genes belonging to the enrichment categories divided by the total number of genes per category.

**Supplementary figure 2:**
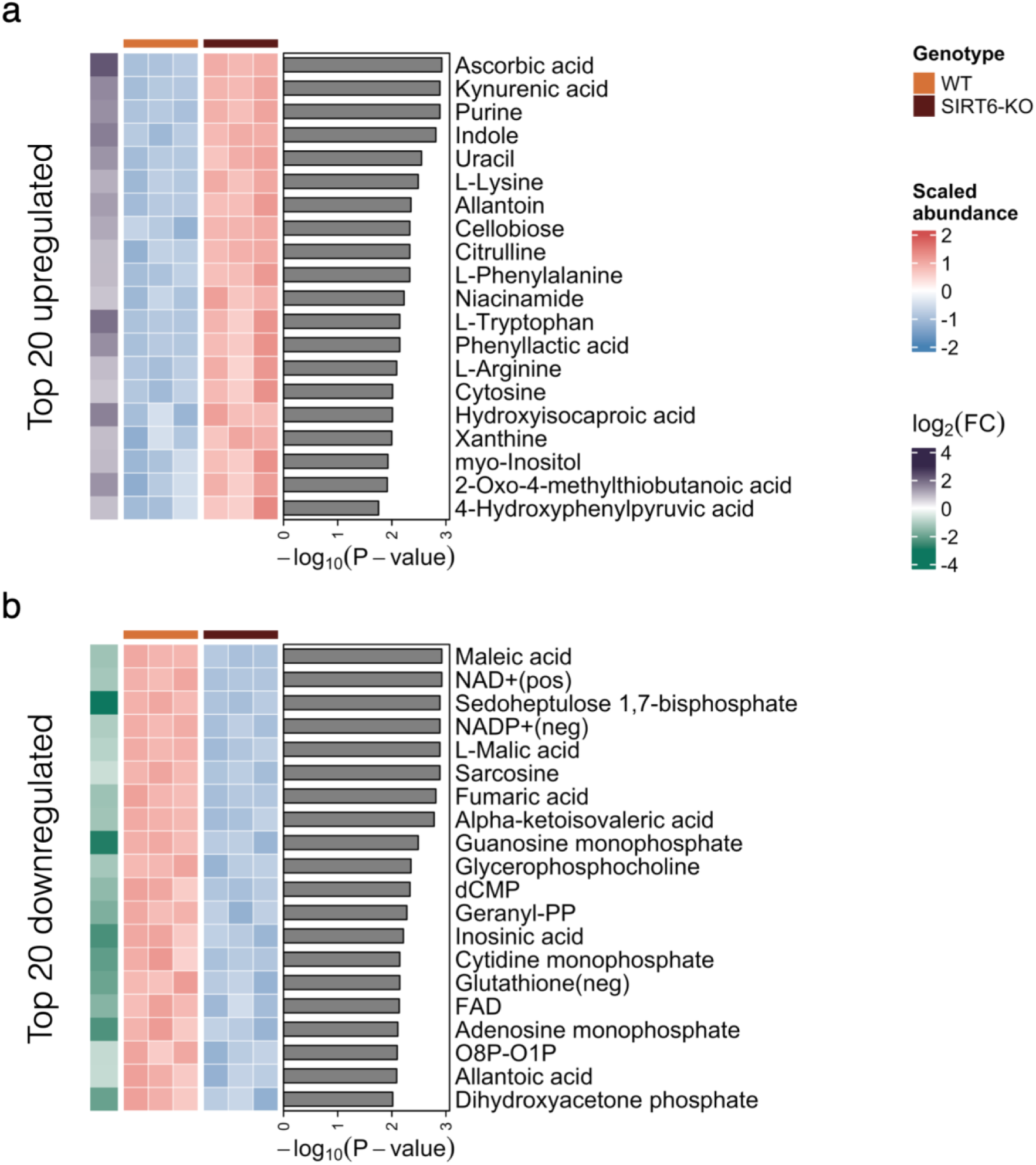
mESC metabolomics (related to Fig. 2). **(a,b)** Heatmaps showing abundances of the top 20 most significantly upregulated (panel a) and downregulated (panel b) in SIRT6-KO (brown) compared to WT (orange) metabolites. Row annotations on the left of the heatmaps represent log_2_ Fold Change values corresponding to the metabolites. Barplots on the right side of the heatmaps represent −log_10_ transformed FDR p-value of the metabolites.

**Supplementary figure 3:**
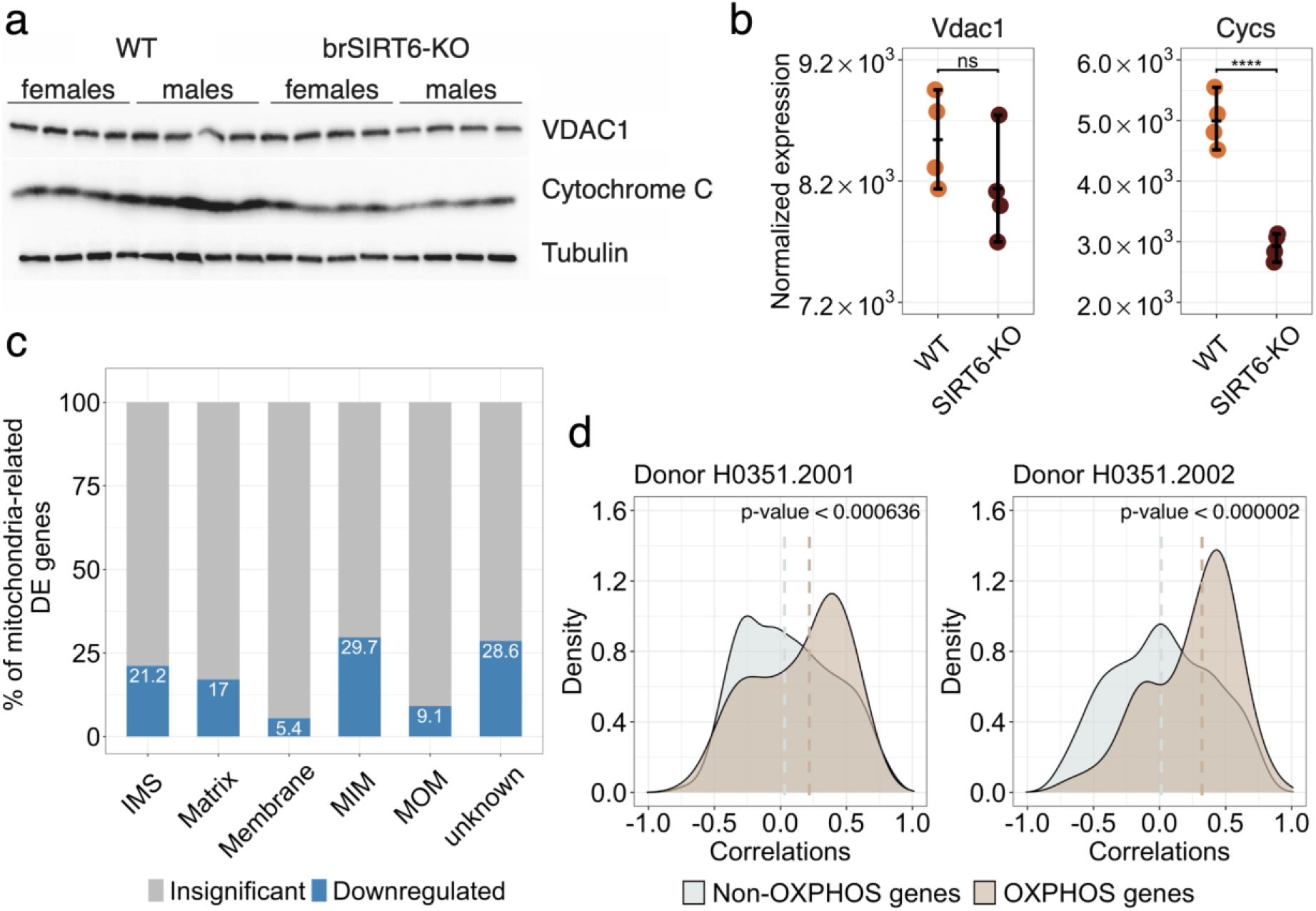
SIRT6 regulates OXPHOS-related genes (related to Fig. 3). **(a)** Western blot analysis of total brain extracts from WT (n=8) and brSIRT6-KO brains (n=8). The total brain fraction was prepared as described in Materials and Methods. Membrane blots were incubated with antibodies against Cytochrome C, Tubulin and VDAC. **(b)** *Vdac1* and *Cycs* expression levels in WT and SIRT6-KO RNA-seq profiles. **(c)** The percentage of significant (blue bars) and insignificant (gray bars) genes across mitochondrial compartments. ‘IMS’ denotes intermembrane space, ‘MIM’ denotes mitochondrial inner membrane, and ‘MOM’ corresponds to the mitochondrial outer membrane. () Spearman’s correlation value distributions for SIRT6 with OXPHOS-related (brown shapes) and other mitochondria-related genes (blue shapes) in the Allen Brain Atlas RNA-seq datasets of two donors (H0351.2001, H0351.2002). Brown and blue dashed lines correspond to medians of correlation distributions for OXPHOS and non-OXPHOS genes, respectively. Statistical significance for SIRT6 correlation with OXPHOS-related genes is calculated via permutation test.

**Supplementary figure 4:**
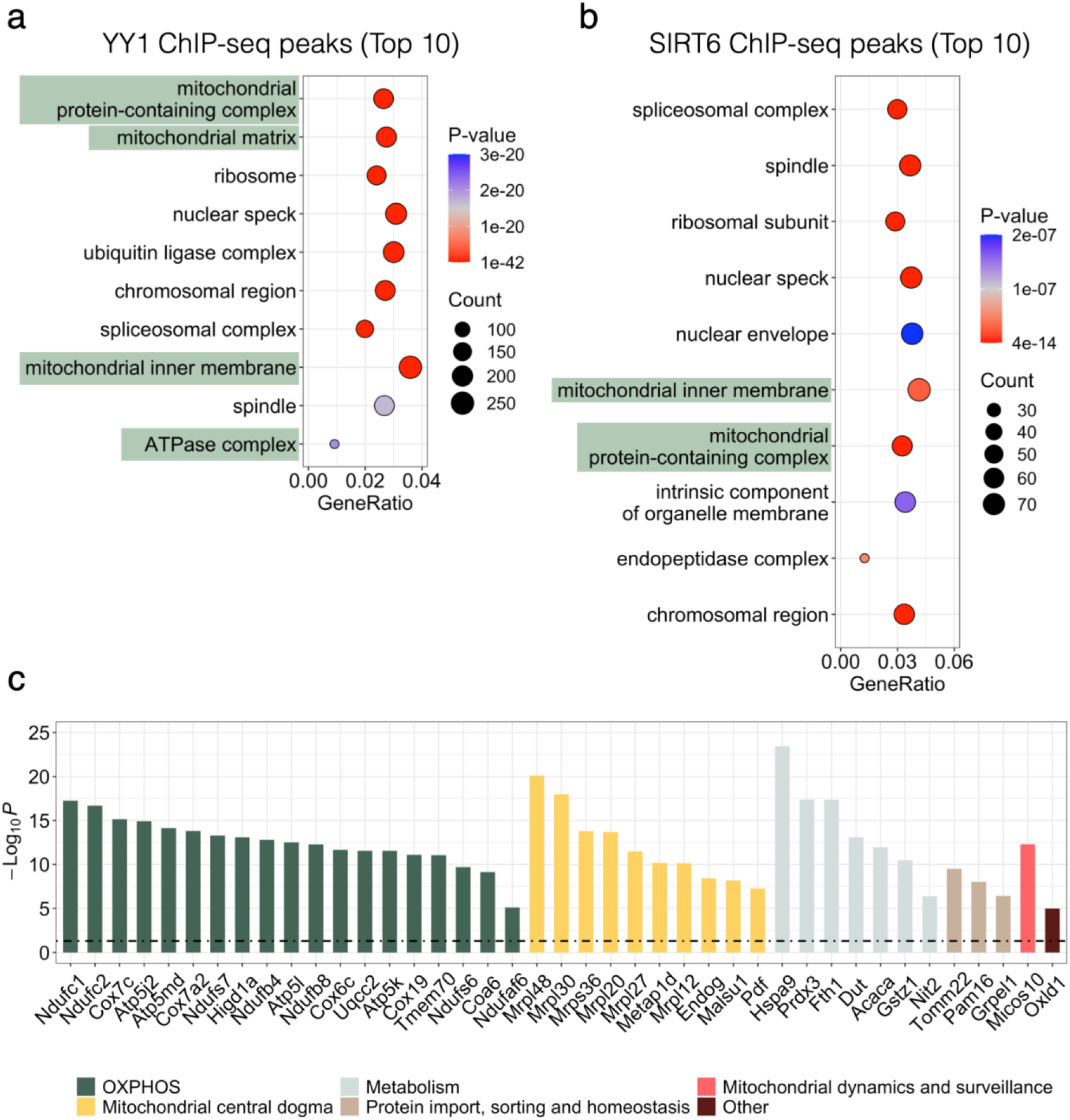
Analysis of public YY1 and SIRT6 ChIP-seq datasets (related to Fig. 4). **(a-b)** Top 10 significant cellular component terms from GO ontology analysis of genes associated with YY1 (panel a) and SIRT6 (panel b) peaks. Mitochondria-related cell compartments are marked by green. **(c)** Barplot showing the expression change magnitudes of genes overlapped between all the datasets presented in Fig. 4d. Bars are colored according to the cellular function of corresponding genes. Black dashed denotes significance cut off for −log_10_ FDR p-value.

**Supplementary figure 5:**
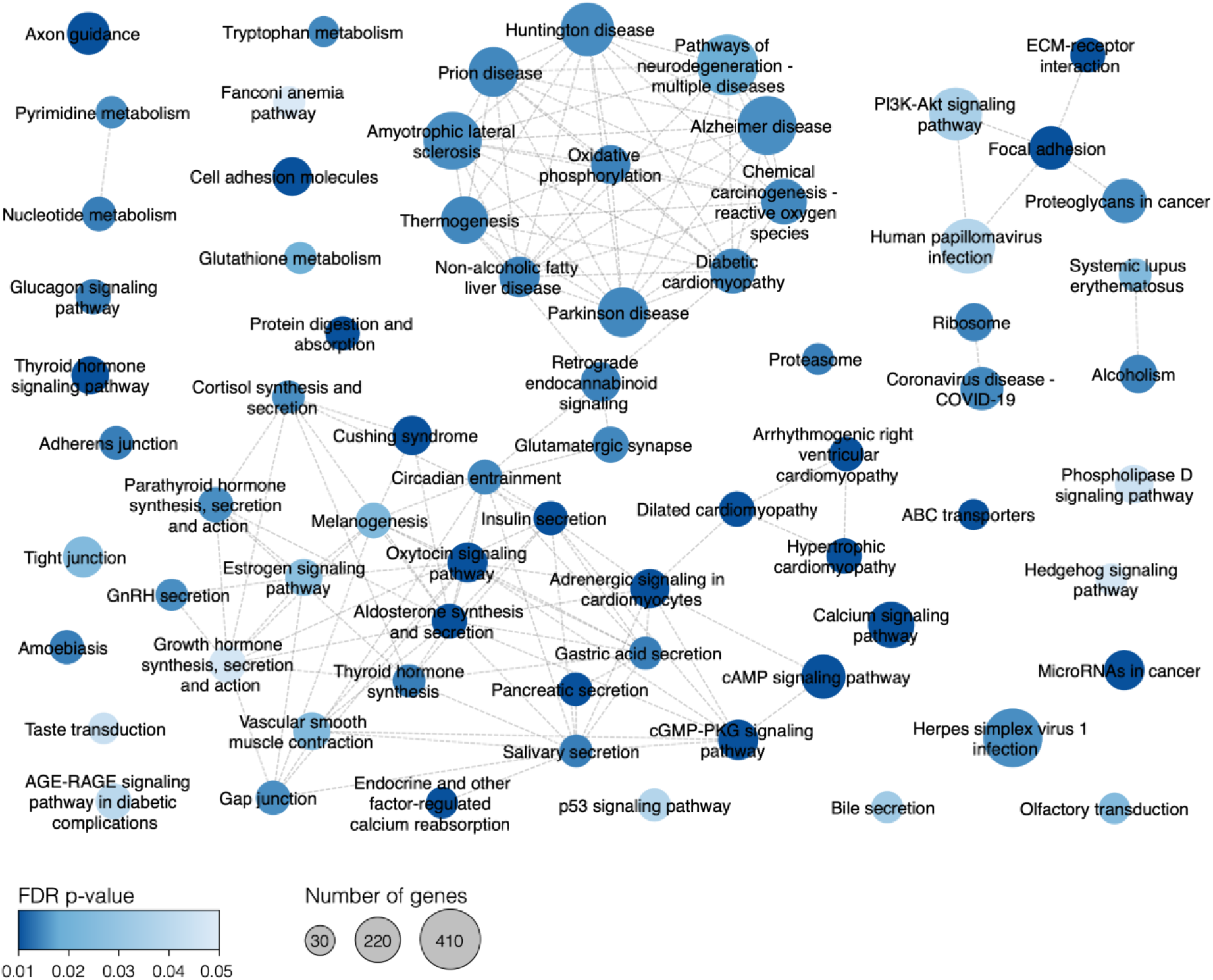
Full network of enriched KEGG pathways (related to Fig. 5). Each circle represents an enriched pathway in GSEA analysis and is colored according to the FDR p-value. The size of circles corresponds to the number of detected genes related to the particular pathway.

## Notes

### Competing Interest Statement

The authors have declared no competing interest.

## References

1 Lopez-Otin, C., Blasco, M. A., Partridge, L., Serrano, M. & Kroemer, G. The hallmarks of aging. Cell 153, 1194–1217, doi:10.1016/j.cell.2013.05.039 (2013).

2 Mattson, M. P. & Arumugam, T. V. Hallmarks of Brain Aging: Adaptive and Pathological Modification by Metabolic States. Cell Metab 27, 1176–1199, doi:10.1016/j.cmet.2018.05.011 (2018).

3 Hernandez-Segura, A., Nehme, J. & Demaria, M. Hallmarks of Cellular Senescence. Trends Cell Biol 28, 436–453, doi:10.1016/j.tcb.2018.02.001 (2018).

4 Sahin, E. & DePinho, R. A. Axis of ageing: telomeres, p53 and mitochondria. Nat Rev Mol Cell Biol 13, 397–404, doi:10.1038/nrm3352 (2012).

5 Peters, R. Ageing and the brain. Postgrad Med J 82, 84–88, doi:10.1136/pgmj.2005.036665 (2006).

6 Blinkouskaya, Y., Cacoilo, A., Gollamudi, T., Jalalian, S. & Weickenmeier, J. Brain aging mechanisms with mechanical manifestations. Mech Ageing Dev 200, 111575, doi:10.1016/j.mad.2021.111575 (2021).

7 Raz, N., Lindenberger, U., Rodrigue, K. M., Kennedy, K. M., Head, D., Williamson, A., Dahle, C., Gerstorf, D. & Acker, J. D. Regional brain changes in aging healthy adults: general trends, individual differences and modifiers. Cereb Cortex 15, 1676–1689, doi:10.1093/cercor/bhi044 (2005).

8 Rine, J., Strathern, J. N., Hicks, J. B. & Herskowitz, I. A suppressor of mating-type locus mutations in Saccharomyces cerevisiae: evidence for and identification of cryptic mating-type loci. Genetics 93, 877–901, doi:10.1093/genetics/93.4.877 (1979).

9 Braunstein, M., Rose, A. B., Holmes, S. G., Allis, C. D. & Broach, J. R. Transcriptional silencing in yeast is associated with reduced nucleosome acetylation. Genes Dev 7, 592–604, doi:10.1101/gad.7.4.592 (1993).

10 Tanner, K. G., Landry, J., Sternglanz, R. & Denu, J. M. Silent information regulator 2 family of NAD-dependent histone/protein deacetylases generates a unique product, 1-O-acetyl-ADP-ribose. Proc Natl Acad Sci U S A 97, 14178–14182, doi:10.1073/pnas.250422697 (2000).

11 Mostoslavsky, R., Chua, K. F., Lombard, D. B., Pang, W. W., Fischer, M. R., Gellon, L., Liu, P., Mostoslavsky, G., Franco, S., Murphy, M. M. et al. Genomic instability and aging-like phenotype in the absence of mammalian SIRT6. Cell 124, 315–329, doi:10.1016/j.cell.2005.11.044 (2006).

12 Van Meter, M., Kashyap, M., Rezazadeh, S., Geneva, A. J., Morello, T. D., Seluanov, A. & Gorbunova, V. SIRT6 represses LINE1 retrotransposons by ribosylating KAP1 but this repression fails with stress and age. Nat Commun 5, 5011, doi:10.1038/ncomms6011 (2014).

13 Onn, L., Portillo, M., Ilic, S., Cleitman, G., Stein, D., Kaluski, S., Shirat, I., Slobodnik, Z., Einav, M., Erdel, F. et al. SIRT6 is a DNA double-strand break sensor. Elife 9, doi:10.7554/eLife.51636 (2020).

14 Toiber, D., Erdel, F., Bouazoune, K., Silberman, D. M., Zhong, L., Mulligan, P., Sebastian, C., Cosentino, C., Martinez-Pastor, B., Giacosa, S. et al. SIRT6 recruits SNF2H to DNA break sites, preventing genomic instability through chromatin remodeling. Mol Cell 51, 454–468, doi:10.1016/j.molcel.2013.06.018 (2013).

15 Mao, Z., Hine, C., Tian, X., Van Meter, M., Au, M., Vaidya, A., Seluanov, A. & Gorbunova, V. SIRT6 promotes DNA repair under stress by activating PARP1. Science 332, 1443–1446, doi:10.1126/science.1202723 (2011).

16 Michishita, E., McCord, R. A., Berber, E., Kioi, M., Padilla-Nash, H., Damian, M., Cheung, P., Kusumoto, R., Kawahara, T. L., Barrett, J. C. et al. SIRT6 is a histone H3 lysine 9 deacetylase that modulates telomeric chromatin. Nature 452, 492–496, doi:10.1038/nature06736 (2008).

17 Roichman, A., Elhanati, S., Aon, M. A., Abramovich, I., Di Francesco, A., Shahar, Y., Avivi, M. Y., Shurgi, M., Rubinstein, A., Wiesner, Y. et al. Restoration of energy homeostasis by SIRT6 extends healthy lifespan. Nat Commun 12, 3208, doi:10.1038/s41467-021-23545-7 (2021).

18 Kaluski, S., Portillo, M., Besnard, A., Stein, D., Einav, M., Zhong, L., Ueberham, U., Arendt, T., Mostoslavsky, R., Sahay, A. et al. Neuroprotective Functions for the Histone Deacetylase SIRT6. Cell Rep 18, 3052–3062, doi:10.1016/j.celrep.2017.03.008 (2017).

19 Li, X., Liu, L., Li, T., Liu, M., Wang, Y., Ma, H., Mu, N. & Wang, H. SIRT6 in Senescence and Aging-Related Cardiovascular Diseases. Front Cell Dev Biol 9, 641315, doi:10.3389/fcell.2021.641315 (2021).

20 Khan, R. I., Nirzhor, S. S. R. & Akter, R. A Review of the Recent Advances Made with SIRT6 and its Implications on Aging Related Processes, Major Human Diseases, and Possible Therapeutic Targets. Biomolecules 8, doi:10.3390/biom8030044 (2018).

21 Garcia-Venzor, A. & Toiber, D. SIRT6 Through the Brain Evolution, Development, and Aging. Front Aging Neurosci 13, 747989, doi:10.3389/fnagi.2021.747989 (2021).

22 Lee, O. H., Kim, J., Kim, J. M., Lee, H., Kim, E. H., Bae, S. K., Choi, Y., Nam, H. S. & Heo, J. H. Decreased expression of sirtuin 6 is associated with release of high mobility group box-1 after cerebral ischemia. Biochem Biophys Res Commun 438, 388–394, doi:10.1016/j.bbrc.2013.07.085 (2013).

23 Portillo, M., Eremenko, E., Kaluski, S., Garcia-Venzor, A., Onn, L., Stein, D., Slobodnik, Z., Zaretsky, A., Ueberham, U., Einav, M. et al. SIRT6-CBP-dependent nuclear Tau accumulation and its role in protein synthesis. Cell Rep 35, 109035, doi:10.1016/j.celrep.2021.109035 (2021).

24 Stein, D., Mizrahi, A., Golova, A., Saretzky, A., Venzor, A. G., Slobodnik, Z., Kaluski, S., Einav, M., Khrameeva, E. & Toiber, D. Aging and pathological aging signatures of the brain: through the focusing lens of SIRT6. Aging (Albany NY) 13, 6420–6441, doi:10.18632/aging.202755 (2021).

25 Mecocci, P., MacGarvey, U., Kaufman, A. E., Koontz, D., Shoffner, J. M., Wallace, D. C. & Beal, M. F. Oxidative damage to mitochondrial DNA shows marked age-dependent increases in human brain. Ann Neurol 34, 609–616, doi:10.1002/ana.410340416 (1993).

26 Wallace, D. C. A mitochondrial paradigm of metabolic and degenerative diseases, aging, and cancer: a dawn for evolutionary medicine. Annu Rev Genet 39, 359–407, doi:10.1146/annurev.genet.39.110304.095751 (2005).

27 Park, C. B. & Larsson, N. G. Mitochondrial DNA mutations in disease and aging. J Cell Biol 193, 809–818, doi:10.1083/jcb.201010024 (2011).

28 Harman, D. Aging: a theory based on free radical and radiation chemistry. J Gerontol 11, 298–300, doi:10.1093/geronj/11.3.298 (1956).

29 Chakrabarti, S., Munshi, S., Banerjee, K., Thakurta, I. G., Sinha, M. & Bagh, M. B. Mitochondrial Dysfunction during Brain Aging: Role of Oxidative Stress and Modulation by Antioxidant Supplementation. Aging Dis 2, 242–256 (2011).

30 Raichle, M. E. & Gusnard, D. A. Appraising the brain’s energy budget. Proc Natl Acad Sci U S A 99, 10237–10239, doi:10.1073/pnas.172399499 (2002).

31 Storozhuk, M. V., Ivanova, S. Y., Balaban, P. M. & Kostyuk, P. G. Possible role of mitochondria in posttetanic potentiation of GABAergic synaptic transmission in rat neocortical cell cultures. Synapse 58, 45–52, doi:10.1002/syn.20186 (2005).

32 Motori, E., Atanassov, I., Kochan, S. M. V., Folz-Donahue, K., Sakthivelu, V., Giavalisco, P., Toni, N., Puyal, J. & Larsson, N. G. Neuronal metabolic rewiring promotes resilience to neurodegeneration caused by mitochondrial dysfunction. Sci Adv 6, eaba8271, doi:10.1126/sciadv.aba8271 (2020).

33 Cunningham, J. T., Rodgers, J. T., Arlow, D. H., Vazquez, F., Mootha, V. K. & Puigserver, P. mTOR controls mitochondrial oxidative function through a YY1-PGC-1alpha transcriptional complex. Nature 450, 736–740, doi:10.1038/nature06322 (2007).

34 Ansari, A., Rahman, M. S., Saha, S. K., Saikot, F. K., Deep, A. & Kim, K. H. Function of the SIRT3 mitochondrial deacetylase in cellular physiology, cancer, and neurodegenerative disease. Aging Cell 16, 4–16, doi:10.1111/acel.12538 (2017).

35 Dai, S. H., Chen, T., Wang, Y. H., Zhu, J., Luo, P., Rao, W., Yang, Y. F., Fei, Z. & Jiang, X. F. Sirt3 protects cortical neurons against oxidative stress via regulating mitochondrial Ca2+ and mitochondrial biogenesis. Int J Mol Sci 15, 14591–14609, doi:10.3390/ijms150814591 (2014).

36 Rath, S., Sharma, R., Gupta, R., Ast, T., Chan, C., Durham, T. J., Goodman, R. P., Grabarek, Z., Haas, M. E., Hung, W. H. W. et al. MitoCarta3.0: an updated mitochondrial proteome now with suborganelle localization and pathway annotations. Nucleic Acids Res 49, D1541–D1547, doi:10.1093/nar/gkaa1011 (2021).

37 Sunkin, S. M., Ng, L., Lau, C., Dolbeare, T., Gilbert, T. L., Thompson, C. L., Hawrylycz, M. & Dang, C. Allen Brain Atlas: an integrated spatio-temporal portal for exploring the central nervous system. Nucleic Acids Res 41, D996–D1008, doi:10.1093/nar/gks1042 (2013).

38 van de Ven, R. A. H., Santos, D. & Haigis, M. C. Mitochondrial Sirtuins and Molecular Mechanisms of Aging. Trends Mol Med 23, 320–331, doi:10.1016/j.molmed.2017.02.005 (2017).

39 Zhang, Y., Chen, K., Sloan, S. A., Bennett, M. L., Scholze, A. R., O’Keeffe, S., Phatnani, H. P., Guarnieri, P., Caneda, C., Ruderisch, N. et al. An RNA-sequencing transcriptome and splicing database of glia, neurons, and vascular cells of the cerebral cortex. J Neurosci 34, 11929–11947, doi:10.1523/JNEUROSCI.1860-14.2014 (2014).

40 Mariottini, C., Scartabelli, T., Bongers, G., Arrigucci, S., Nosi, D., Leurs, R., Chiarugi, A., Blandina, P., Pellegrini-Giampietro, D. E. & Passani, M. B. Activation of the histaminergic H3 receptor induces phosphorylation of the Akt/GSK-3 beta pathway in cultured cortical neurons and protects against neurotoxic insults. J Neurochem 110, 1469–1478, doi:10.1111/j.1471-4159.2009.06249.x (2009).

41 Shao, J., Yang, X., Liu, T., Zhang, T., Xie, Q. R. & Xia, W. Autophagy induction by SIRT6 is involved in oxidative stress-induced neuronal damage. Protein Cell 7, 281–290, doi:10.1007/s13238-016-0257-6 (2016).

42 Rayaprolu, S., Bitarafan, S., Santiago, J. V., Betarbet, R., Sunna, S., Cheng, L., Xiao, H., Nelson, R. S., Kumar, P., Bagchi, P. et al. Cell type-specific biotin labeling in vivo resolves regional neuronal and astrocyte proteomic differences in mouse brain. Nat Commun 13, 2927, doi:10.1038/s41467-022-30623-x (2022).

43 Oberdoerffer, P., Michan, S., McVay, M., Mostoslavsky, R., Vann, J., Park, S. K., Hartlerode, A., Stegmuller, J., Hafner, A., Loerch, P. et al. SIRT1 redistribution on chromatin promotes genomic stability but alters gene expression during aging. Cell 135, 907–918, doi:10.1016/j.cell.2008.10.025 (2008).

44 Wang, X., Patel, N. D., Hui, D., Pal, R., Hafez, M. M., Sayed-Ahmed, M. M., Al-Yahya, A. A. & Michaelis, E. K. Gene expression patterns in the hippocampus during the development and aging of Glud1 (Glutamate Dehydrogenase 1) transgenic and wild type mice. BMC Neurosci 15, 37, doi:10.1186/1471-2202-15-37 (2014).

45 Chabi, B., Ljubicic, V., Menzies, K. J., Huang, J. H., Saleem, A. & Hood, D. A. Mitochondrial function and apoptotic susceptibility in aging skeletal muscle. Aging Cell 7, 2–12, doi:10.1111/j.1474-9726.2007.00347.x (2008).

46 Jung, E. S., Choi, H., Song, H., Hwang, Y. J., Kim, A., Ryu, H. & Mook-Jung, I. p53-dependent SIRT6 expression protects Abeta42-induced DNA damage. Sci Rep 6, 25628, doi:10.1038/srep25628 (2016).

47 Camacho-Pereira, J., Tarrago, M. G., Chini, C. C. S., Nin, V., Escande, C., Warner, G. M., Puranik, A. S., Schoon, R. A., Reid, J. M., Galina, A. et al. CD38 Dictates Age-Related NAD Decline and Mitochondrial Dysfunction through an SIRT3-Dependent Mechanism. Cell Metab 23, 1127–1139, doi:10.1016/j.cmet.2016.05.006 (2016).

48 Mouchiroud, L., Houtkooper, R. H., Moullan, N., Katsyuba, E., Ryu, D., Canto, C., Mottis, A., Jo, Y. S., Viswanathan, M., Schoonjans, K. et al. The NAD(+)/Sirtuin Pathway Modulates Longevity through Activation of Mitochondrial UPR and FOXO Signaling. Cell 154, 430–441, doi:10.1016/j.cell.2013.06.016 (2013).

49 Imai, S. & Guarente, L. NAD+ and sirtuins in aging and disease. Trends Cell Biol 24, 464–471, doi:10.1016/j.tcb.2014.04.002 (2014).

50 Wood, J. G., Schwer, B., Wickremesinghe, P. C., Hartnett, D. A., Burhenn, L., Garcia, M., Li, M., Verdin, E. & Helfand, S. L. Sirt4 is a mitochondrial regulator of metabolism and lifespan in Drosophila melanogaster. Proc Natl Acad Sci U S A 115, 1564–1569, doi:10.1073/pnas.1720673115 (2018).

51 Min, Z., Gao, J. & Yu, Y. The Roles of Mitochondrial SIRT4 in Cellular Metabolism. Front Endocrinol (Lausanne) 9, 783, doi:10.3389/fendo.2018.00783 (2018).

52 Kincaid, B. & Bossy-Wetzel, E. Forever young: SIRT3 a shield against mitochondrial meltdown, aging, and neurodegeneration. Front Aging Neurosci 5, 48, doi:10.3389/fnagi.2013.00048 (2013).

53 Boxer, L. D., Renthal, W., Greben, A. W., Whitwam, T., Silberfeld, A., Stroud, H., Li, E., Yang, M. G., Kinde, B., Griffith, E. C. et al. MeCP2 Represses the Rate of Transcriptional Initiation of Highly Methylated Long Genes. Mol Cell 77, 294–309 e299, doi:10.1016/j.molcel.2019.10.032 (2020).

54 Sebastian, C., Zwaans, B. M., Silberman, D. M., Gymrek, M., Goren, A., Zhong, L., Ram, O., Truelove, J., Guimaraes, A. R., Toiber, D. et al. The histone deacetylase SIRT6 is a tumor suppressor that controls cancer metabolism. Cell 151, 1185–1199, doi:10.1016/j.cell.2012.10.047 (2012).

55 Chen, S., Zhou, Y., Chen, Y. & Gu, J. fastp: an ultra-fast all-in-one FASTQ preprocessor. Bioinformatics 34, i884–i890, doi:10.1093/bioinformatics/bty560 (2018).

56 Dobin, A., Davis, C. A., Schlesinger, F., Drenkow, J., Zaleski, C., Jha, S., Batut, P., Chaisson, M. & Gingeras, T. R. STAR: ultrafast universal RNA-seq aligner. Bioinformatics 29, 15–21, doi:10.1093/bioinformatics/bts635 (2013).

57 Patro, R., Duggal, G., Love, M. I., Irizarry, R. A. & Kingsford, C. Salmon provides fast and bias-aware quantification of transcript expression. Nat Methods 14, 417–419, doi:10.1038/nmeth.4197 (2017).

58 Love, M. I., Huber, W. & Anders, S. Moderated estimation of fold change and dispersion for RNA-seq data with DESeq2. Genome Biol 15, 550, doi:10.1186/s13059-014-0550-8 (2014).

59 Yu, G., Wang, L. G., Han, Y. & He, Q. Y. clusterProfiler: an R package for comparing biological themes among gene clusters. OMICS 16, 284–287, doi:10.1089/omi.2011.0118 (2012).

60 Shannon, P., Markiel, A., Ozier, O., Baliga, N. S., Wang, J. T., Ramage, D., Amin, N., Schwikowski, B. & Ideker, T. Cytoscape: a software environment for integrated models of biomolecular interaction networks. Genome Res 13, 2498–2504, doi:10.1101/gr.1239303 (2003).

61 Pang, Z., Chong, J., Zhou, G., de Lima Morais, D. A., Chang, L., Barrette, M., Gauthier, C., Jacques, P. E., Li, S. & Xia, J. MetaboAnalyst 5.0: narrowing the gap between raw spectra and functional insights. Nucleic Acids Res 49, W388–W396, doi:10.1093/nar/gkab382 (2021).

62 Kanehisa, M., Sato, Y., Kawashima, M., Furumichi, M. & Tanabe, M. KEGG as a reference resource for gene and protein annotation. Nucleic Acids Res 44, D457–462, doi:10.1093/nar/gkv1070 (2016).

63 Gu, Z., Eils, R. & Schlesner, M. Complex heatmaps reveal patterns and correlations in multidimensional genomic data. Bioinformatics 32, 2847–2849, doi:10.1093/bioinformatics/btw313 (2016).

64 Etchegaray, J. P., Chavez, L., Huang, Y., Ross, K. N., Choi, J., Martinez-Pastor, B., Walsh, R. M., Sommer, C. A., Lienhard, M., Gladden, A. et al. The histone deacetylase SIRT6 controls embryonic stem cell fate via TET-mediated production of 5-hydroxymethylcytosine. Nat Cell Biol 17, 545–557, doi:10.1038/ncb3147 (2015).

65 Oki, S., Ohta, T., Shioi, G., Hatanaka, H., Ogasawara, O., Okuda, Y., Kawaji, H., Nakaki, R., Sese, J. & Meno, C. ChIP-Atlas: a data-mining suite powered by full integration of public ChIP-seq data. EMBO Rep 19, doi:10.15252/embr.201846255 (2018).

66 Yu, G., Wang, L. G. & He, Q. Y. ChIPseeker: an R/Bioconductor package for ChIP peak annotation, comparison and visualization. Bioinformatics 31, 2382–2383, doi:10.1093/bioinformatics/btv145 (2015).

67 Gel, B. & Serra, E. karyoploteR: an R/Bioconductor package to plot customizable genomes displaying arbitrary data. Bioinformatics 33, 3088–3090, doi:10.1093/bioinformatics/btx346 (2017).

68 Barrett, T., Wilhite, S. E., Ledoux, P., Evangelista, C., Kim, I. F., Tomashevsky, M., Marshall, K. A., Phillippy, K. H., Sherman, P. M., Holko, M. et al. NCBI GEO: archive for functional genomics data sets--update. Nucleic Acids Res 41, D991–995, doi:10.1093/nar/gks1193 (2013).

